# Inter-domain dynamics and fibrinolytic activity of matrix metalloprotease-1

**DOI:** 10.1101/853796

**Authors:** Lokender Kumar, Joan Planas-Iglesias, Chase Harms, Sumaer Kamboj, Derek Wright, Judith Klein-Seetharaman, Susanta K. Sarkar

## Abstract

The roles of protein conformational dynamics and allostery in function are well-known. However, the roles that inter-domain dynamics have in function are not entirely understood. We used matrix metalloprotease-1 (MMP1) as a model system to study the relationship between inter-domain dynamics and activity because MMP1 has diverse substrates. Here we focus on fibrin, the primary component of a blood clot. Water-soluble fibrinogen, following cleavage by thrombin, self-polymerize to form water-insoluble fibrin. We studied the inter-domain dynamics of MMP1 on fibrin without crosslinks using single-molecule Forster Resonance Energy Transfer (smFRET). We observed that the distance between the catalytic and hemopexin domains of MMP1 increases or decreases as the MMP1 activity increases or decreases, respectively. We modulated the activity using 1) an active site mutant (E219Q) of MMP1, 2) MMP9, another member of the MMP family that increases the activity of MMP1, and 3) tetracycline, an inhibitor of MMP1. We fitted the histograms of smFRET values to a sum of two Gaussians and the autocorrelations to an exponential and power law. We modeled the dynamics as a two-state Poisson process and calculated the kinetic rates from the histograms and autocorrelations. Activity-dependent inter-domain dynamics may enable allosteric control of the MMP1 function.

## Introduction

An understanding of the relationship between the structure and dynamics of a protein is crucial for understanding its function^1^. Researchers have shown the correlation between the function and intra-domain dynamics within a protein domain at different hierarchies of timescales^2-4^. However, the roles of inter-domain dynamics in function are not entirely understood^5,6^. Matrix metalloproteases (MMPs) are suitable for investigating the functional relationship between inter-domain dynamics and activity because the sequence of the catalytic domain remains conserved across the 23-member family^7^. The differences in activities among MMPs likely originate from the hemopexin domain with significant variations in the sequence^7^. The hemopexin domain seems to influence substrate/ligand specificity and activation/inhibition of various MMPs^8^. Also, prior research has reported the regulation of MMP1 catalytic activity by the hemopexin domain^9^. MMPs are capable of degrading numerous proteins^10^, including collagen, the primary component of the extracellular matrix (ECM) that provides a scaffold for cells to maintain tissue integrity^11^. MMPs are calcium- and zinc-dependent enzymes that are produced by cells in pro forms, i.e., they need to be activated by cleaving off the pro domain for activity^11,12^. MMP1, a collagenase that degrades triple-helical type-1 collagen, stands out in the 23-member MMP family because it has crystal structures available^13^. Also, MMP1 itself is a broad-spectrum protease, an attribute that we previously used to degrade *E. coli* proteins and purify recombinant MMP1 using a protease-based method^14^. Extensive studies of MMP1 interacting with collagen monomers have revealed significant insights into the MMP1 activity, including the roles of conformational dynamics^9,15-22^. MMP1 consists of a catalytic domain that degrades substrates, a hemopexin domain that helps MMPs bind to the substrates, and a linker that mediates communications between the two domains (**Figure 1**).

**Figure 1.**
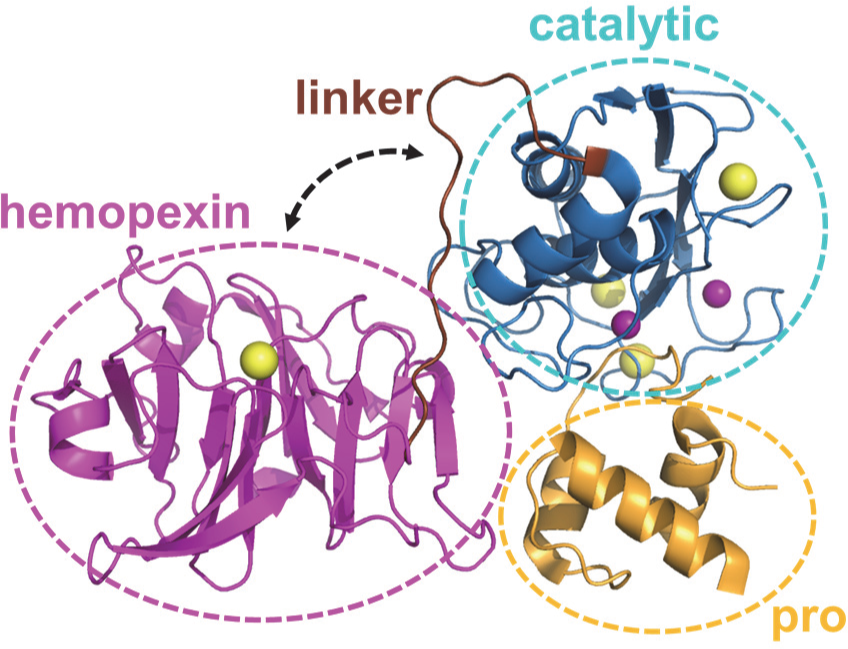
MMP1 structure (PDB ID: ISU3). The hemopexin domain is connected to the catalytic domain by a flexible linker. The pro domain is cleaved off to activate MMPs for catalysis. Yellow and purple spheres represent the van der Waals radii of the calcium and zinc atoms, respectively. The pro domain, the catalytic domain, the linker, and the hemopexin domains are roughly defined by the ranges of residues D32-Q99, F100-Y260, G261-C278, and D279-C466, respectively.

Recently, we showed that inter-domain dynamics of MMP1 on type-1 collagen fibrils correlate with activity, and the two domains can allosterically communicate via the linker region as well as the substrate^23^. Since MMP1 is promiscuous, we investigated whether the insights into collagen fibril degradation applies to other substrates. In this paper, we focus on MMP1 inter-domain dynamics and activity on fibrin, the primary component of a blood clot. Fibrin is a critical physiological substrate because ∼900,000 patients suffer from complications from clot formation leading to ∼300,000 deaths^24^ and ∼$28 billion annual cost^25^ in the US alone. The number of adults with clot-related health issues is estimated to reach ∼1.82 million by 2050^26^. Blood clot formation through the coagulation cascade is a part of the natural response to injury and cut and prevents loss of blood from severed blood vessels^27^. The life-saving process of clot formation can become life-threatening if a clot blocks a blood vessel. Indeed, pathologies of clot formation include heart attack, stroke, and pulmonary embolism, accounting for ∼50% of all hospital deaths.

Reperfusion, the act of restoring blood flow, with tissue plasminogen activator (tPA) remains the standard treatment for ischemic stroke^28^. Plasminogen becomes plasmin after cleavage by tPA and is able to degrade blood clots^29^. However, initiating tPA treatment after 3-4.5 h of stroke onset can lead to adverse side effects, notably, hemorrhagic transformation^30^. Therefore, it is important to explore alternative approaches that could either replace tPA treatment or extend the time window for tPA perfusion beyond the current time frame^28^. For example, staphopain A and staphopain B, cysteine proteases released by pathogenic Gram-positive bacteria *Staphylococcus aureus*, impair clotting by degrading fibrinogen^31^. Also, the observation of clot degradation in plasminogen-deleted animals suggests alternative fibrinolytic pathways^32^. Many proteases other than plasmin have shown fibrinolytic activity. MMPs are potential alternatives in the fibrinolytic system because MMPs are in the blood both in pro^33,34^ and activated^35-38^ forms. There are reports of fibrinolytic activity of MMP3^39^, MMP9^40^, and MMP14^41^, but the field is relatively unexplored^42^. Prior research has reported that MMP1 does not have significant fibrinolytic activity^39^, which is in contrast to our observations presented in this paper.

Here we show that MMP1 inter-domain dynamics on fibrin are activity-dependent and modulated by an enhancer or an inhibitor of MMP activity. We calculated the distance between S142 in the catalytic domain and S366 in the hemopexin domain of MMP1 using smFRET measurements. The area-normalized histograms of smFRET values represent the distributions of conformations, whereas the normalized autocorrelations represent correlations between structures at different time points. From the histograms, we defined two MMP1 structures with 1) inter-domain distance of ∼4.5 nm and 2) inter-domain distance of ∼5.4 nm as the closed and open states, respectively. A comparison of active and active site mutant suggests that the open conformation of MMP1 is functionally essential. The open conformations with well-separated domains appear more in the presence of an activity enhancer (MMP9) and less in the presence of an inhibitor (tetracycline). From autocorrelations, we learned that inter-domain dynamics are not entirely random and have exponentially-distributed correlations. We fitted a sum of two Gaussians to the histograms and an exponential to the autocorrelations. We modeled the dynamics as a two-state process, where the locations of two states are the centers of Gaussian fits, and the sum of two kinetic rates between the two states is the decay rate of exponential fits. Anisotropic Network Modeling (ANM) of MMP1 dynamics revealed that a larger separation between the two domains (open conformation) often accompanies a larger catalytic pocket opening between N171 and T230. A larger catalytic pocket opening, in turn, enables closer proximity to the three chains of fibrin. ANM simulations further revealed that the α-chain of fibrin is closest to the MMP1 active site, whereas the γ-chain is furthest. SDS PAGE of ensemble fibrin degradation confirmed that MMP1 cleaves the α-chain first, followed by the β- and γ-chains. The synergistic combination of smFRET, stochastic simulations, ANM simulations, and ensemble assays provides an integrative approach to investigate the functional roles of inter-domain dynamics. This approach applies to other biochemical processes where water-soluble enzymes interact with water-insoluble substrates.

## Results and discussion

### Single-molecule measurement of MMP1 inter-domain dynamics on fibrin without crosslinks

Water-soluble fibrinogen, upon limited cleavage by thrombin, self-polymerize into water-insoluble fibrin without crosslinks. The addition of factor XIII and CaCl_2_ creates fibrin with crosslinks. For single molecule measurements, we used fibrin without crosslinks because we used a crystal structure of fibrin without crosslinks for simulations. Besides, fibrin without crosslinks is optically clearer than fibrin with crosslinks, which facilitates imaging with less background. To measure MMP1 inter-domain dynamics at the single molecule level, we measured smFRET between two dyes attached to the catalytic and hemopexin domains of MMP1. We mutated S142 in the catalytic domain and S366 in the hemopexin domain of MMP1 to cysteines for labeling. The distances between S142 and S366 are ∼5.4 nm and ∼4.5 nm for the open and closed conformations of MMP1, respectively. We selected a pair of dyes, Alexa555 and Alexa647, because the Forster radius between the two fluorophores is 5.1 nm. The FRET efficiency is 50% when the distance between the fluorophores is equal to the Forster radius. More importantly, FRET is most sensitive to any distance change around the Forster radius.

Single molecule measurements suffer^43^ from labeling^44^, solution conditions^45^, and properties of fluorophores^46-49^. The stochastic nature of labeling CYS142 and CYS366 can also be problem, which we discussed in our previous publication^23^. Since active MMP1 and active site mutant of MMP1 would be affected equally by these complications, we can distinguish the effects due to activity. Note that the amino acid E219^14^ is the same as E200^13^, which differs because we included 19 residues of the pro domain. In a previous publication, we showed that the specific activities of labeled and unlabeled MMP1 were not affected by labeling^50^. A flexible linker connects the catalytic and hemopexin domains of MMP1 (**Figure 2A**). **Figure 2B** shows fibrin without crosslinks. We measured MMP1 inter-domain dynamics on a thin layer of water-insoluble fibrin in a flow cell using a Total Internal Reflection Fluorescence (TIRF) microscope (**Figure 2C**). **Figure 2D** shows one smFRET trajectory. For each condition, we collected more than 300,000 smFRET values for analysis of MMP1 inter-domain dynamics on fibrin (**Figure 3**).

**Figure 2.**
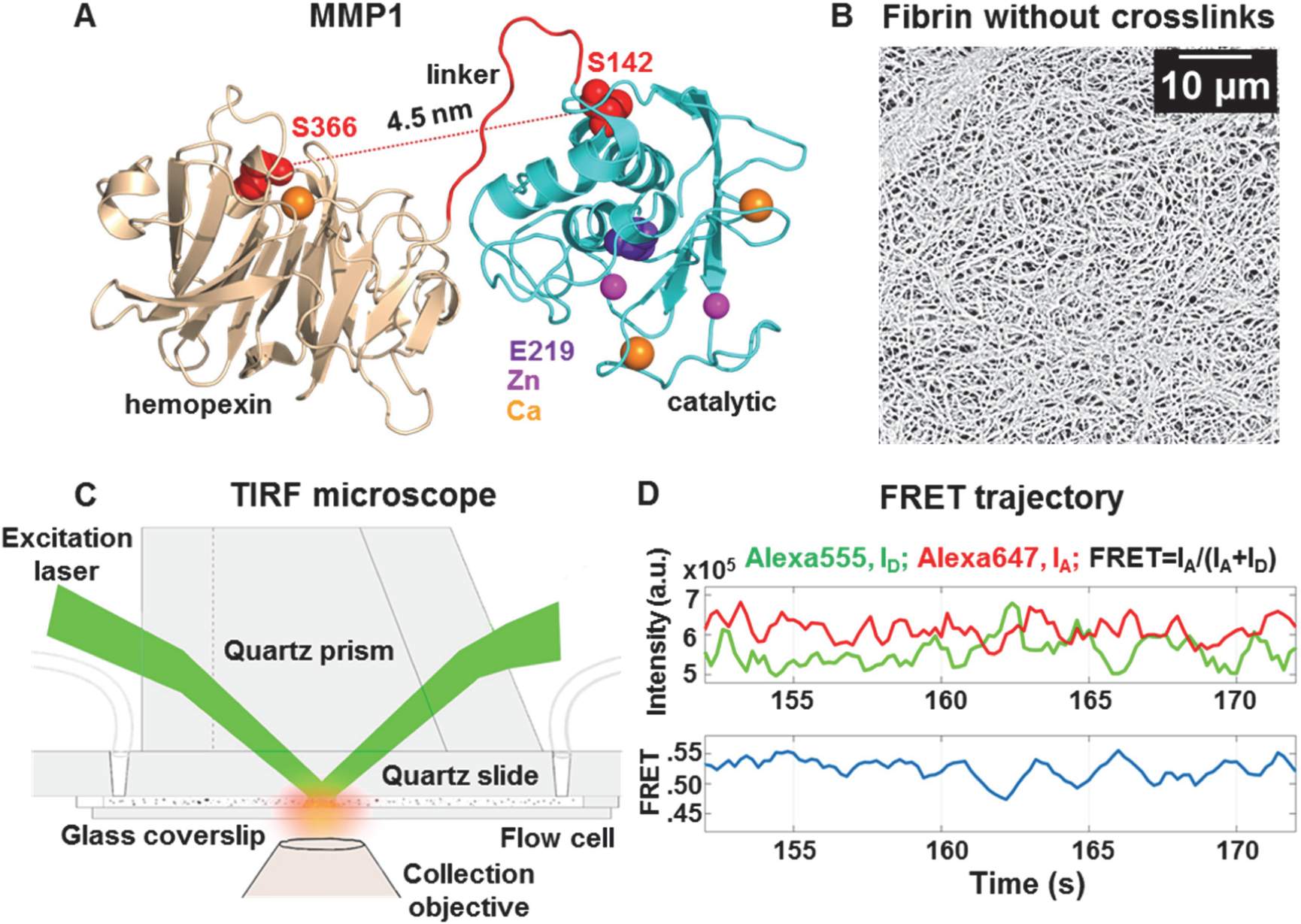
Single-molecule measurement of MMP1 dynamics on fibrin without crosslinks. (**A**) Crystal structure of MMP1 (PDB ID 1SU3). Mutations of S142 and S366 to cysteines enables attaching Alexa555 and Alexa647 dyes. (**B**) Scanning Electron Microscope (SEM) images of fibrin without crosslinks. (**C**) Schematics of the TIRF microscope used for measuring MMP1 inter-domain dynamics on fibrin. (**D**) Emission intensities of the two dyes. (top panel) Low FRET conformations lead to high Alexa555 emission, whereas High FRET conformations lead to Low Alexa555 emission. Anticorrelated Alexa647 and Alexa555 emissions, I_A_ and I_D_, respectively; (bottom panel) Calculated smFRET trajectory to show MMP1 inter-domain dynamics as a function of time.

**Figure 3.**
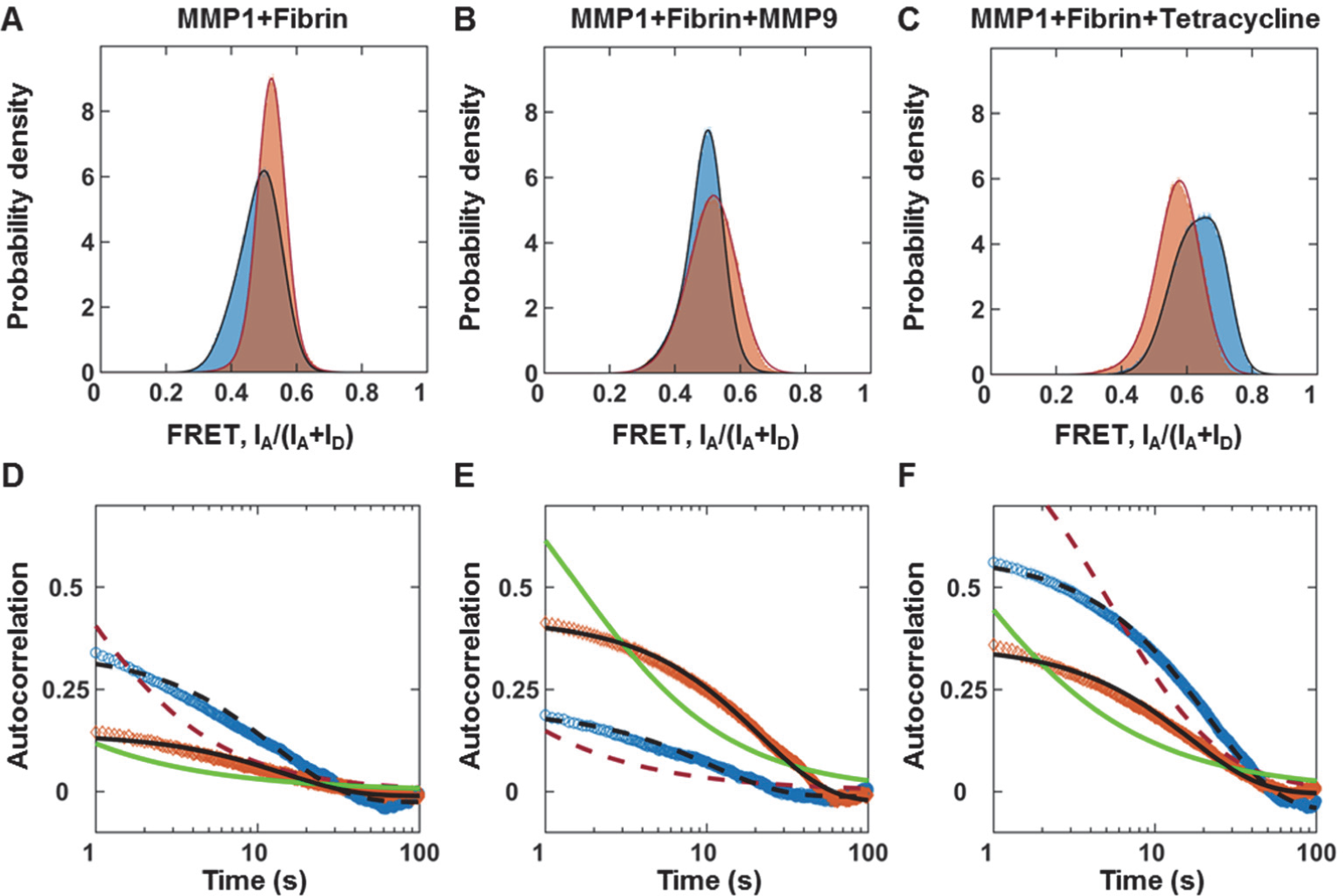
Inter-domain dynamics of MMP1 on fibrin correlates with activity. More than 300,000 smFRET values at 22° C with a 100 ms time resolution for each condition are used to create area-normalized histograms of MMP1 inter-domain distance (bin size=0.005). (**A**) Histograms without ligand, (**B**) Histograms in the presence of MMP9 (an enhancer), and (**C**) Histograms in the presence of tetracycline (an inhibitor) for active (blue) and active site mutant (orange) MMP1. Histograms are fitted to a sum of two Gaussians (active: solid blue line; active site mutant: solid red line). Autocorrelations of MMP1 inter-domain distance are calculated from the time series of smFRET values. (**D**) Autocorrelations without ligand, (**E**) Autocorrelations in the presence of MMP9, and (**F**) Autocorrelations in the presence of tetracycline for active MMP1 (blue) and active site mutant of MMP1 (orange). Autocorrelations are fitted to exponentials and power laws (exponential fit to active: dashed black line; power law fit to active: dashed red line; exponential fit to active site mutant: solid black line; power law fit to active site mutant: solid green line). The error bars in the histograms and autocorrelations represent the square roots of the bin counts and the standard errors of the mean (sem) and are too small to be seen. The supplementary information contains the fit equations and the best-fit parameters for histograms and autocorrelations (**Table S1**). We approximated smFRET efficiency by I_A_/(I_A_+ I_D_)^51^ and has no unit.

### MMP1 inter-domain dynamics on fibrin correlate with activity

Each value of smFRET represents a conformation of MMP1. The histograms of smFRET values measuring inter-domain distances (**Figure 3A**) for active MMP1 and active site mutant of MMP1 suggest that low FRET conformations, where the two domains are well-separated, occur more for active MMP1 (blue histogram) compared to active site mutant (orange histogram). The prevalence of low FRET conformations for active MMP1 suggests that the two domains need to move away from each other for performing catalytic activity on fibrin. Previously, we observed the same effects for MMP1 function on type-1 collagen^23^. Collagen degradation by MMP1 is enhanced by MMP9 and inhibited by tetracycline. For collagen, we found that low FRET conformations appear more for MMP9 and less for tetracycline consistent with the enhancement and inhibition of MMP1 activity. To test whether or not MMP1 follows similar inter-domain dynamics when bound to fibrin, we performed smFRET measurements in the presence of MMP9 and tetracycline. In the presence of tetracycline, low FRET conformations of MMP1 on fibrin significantly disappear on fibrin (**Figure 3C**), as observed on type-1 collagen. Surprisingly, in the presence of MMP9, the MMP1 inter-domain dynamics on fibrin (**Figure 3B**) show more high FRET states in contrast to the occurrence of more low FRET states on collagen. To determine how a conformation at one time point correlates with a conformation at another time point, we calculated the autocorrelations of conformations (**Figure 3D-F**). Without any ligand, the correlation of dynamics on fibrin (**Figure 3D**) is higher for active MMP1 at shorter times, similar to previously reported dynamics on collagen. However, the correlations in the presence of MMP9 and tetracycline show opposite orders on fibrin (**Figure 3E** and **3F**) and collagen (previously reported)^23^. The correlated motions indicate a decrease in conformational entropy. They can affect kinetics and thermodynamics of biological processes, including catalysis^52^. The observed modulations of correlations under different conditions suggest allosterically controlled inter-domain communications. The quantitative comparisons of best-fit parameters (**Table S1**) strongly indicate that inter-domain dynamics and activity of MMP1 are functionally related and allosteric. Further, a comparison with inter-domain dynamics on fibrin (this paper) and collagen^23^ suggest that MMP1 undergoes substrate-dependent and allosterically-controlled dynamics.

### A two-state Poisson process describes MMP1 inter-domain dynamics on fibrin

While the histograms reveal distributions of conformations, the autocorrelations reveal whether or not conformations at different time points are related. We found that a sum of two Gaussians fits the histograms of smFRET values. Power law fits the autocorrelations for two-state simulations without noise at millisecond timescales and molecular dynamics simulations at picosecond timescales for collagen^23^. As such, we tried to fit both power law and exponential to the autocorrelations for fibrin. The fit equations and best-fit parameters are in the supplementary information. The exponential fits to autocorrelations enable a more straightforward interpretation of the decay rates of correlations if we approximate the conformations of MMP1 as a two-state Poisson process^23^. As a result, we can establish a quantitative connection between the histograms and autocorrelations to calculate the kinetic rates between the two states. To test this, we considered the two centers of Gaussian fits to histograms for MMP1 without ligands (**Figure 3A**) as the two states, S1 (low FRET) and S2 (high FRET). We defined the two kinetic rates as k1 (S1→S2) and k2 (S2→S1) for interconversion between S1 and S2. For a two-state system, we calculated k1/k2 from the ratio of Gaussian area(S2)/area(S1) and k1+k2 from the decay rates of autocorrelations. We solved the two equations to calculate k1 and k2 (**Table S1C**) for both active MMP1 and active site mutant of MMP1. With experimentally-determined S1, S2, k1, k2, and noise (widths of the histograms), we simulated smFRET trajectories for active MMP1 and active site mutant of MMP1 and analyzed the same way as we did experimental smFRET trajectories (**Figure 4**). For comparison, experimentally-determined inputs and recovered parameters from two-state simulations are in **Table S2**. In summary, we show that a two-state system describes MMP1 inter-domain dynamics.

**FIGURE 4.**
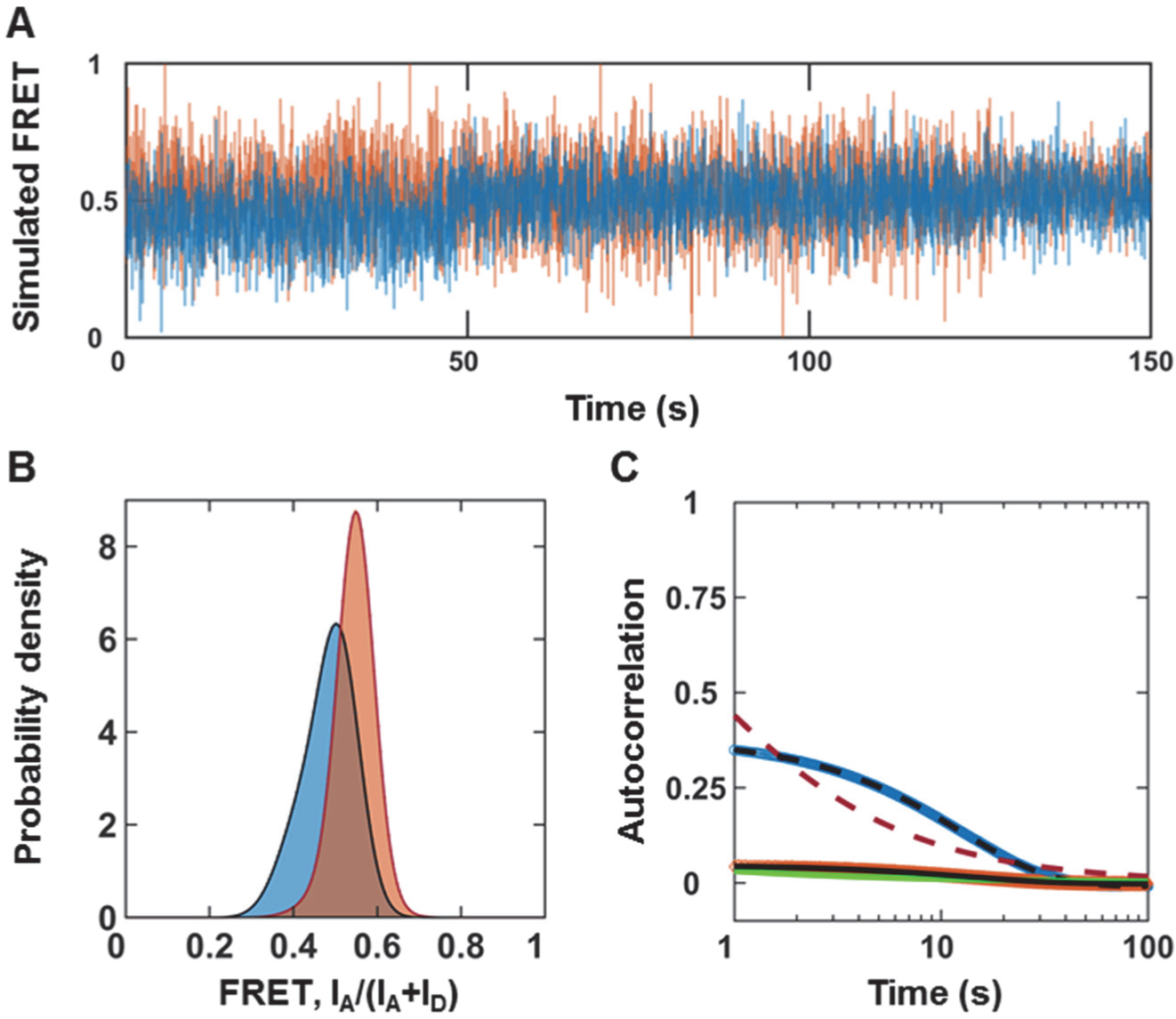
MMP1 inter-domain dynamics as a two-state system. (**A**) Examples of simulated smFRET trajectories with noise for active MMP1 (blue) and active site mutant of MMP1 (orange) using experimentally-determined parameters for MMP1 without tetracycline. (**B**) Area-normalized histograms of simulated smFRET values with best fits to a sum of two Gaussians (solid black line). (**C**) Autocorrelations of simulated smFRET trajectories with best fits to exponentials (active: dashed black line; active site mutant: solid black line). As expected, power law did not fit autocorrelations (active: dashed red line; active site mutant: solid green line). k1+k2 was recovered from exponential fits with and without noise. The error bars are the standard errors of mean for histograms and autocorrelations and are too small to be seen.

For active MMP1 without ligands, the two states are S1=0.42 and S2=0.51 on fibrin (**Table S1A**), which are similar to the two states S1=0.44 and S2=0.55 on collagen. However, the correlation decay rate of 0.08 s^−1^ on fibrin (**Table S1C**) is lower than the rate of 0.13 s^−1^ on collagen. In the presence of MMP9 and tetracycline, the two states and kinetic rates change even though a two-state description still applies for active MMP1 and active site mutant of MMP1 on fibrin (**Figure 3** and **Table S1**) and collagen. In other words, MMP1 undergoes substrate-dependent inter-domain dynamics that follow a two-state Poisson process.

The two-state description also reveals the importance of noise in autocorrelations. Without noise, both power law and exponential fit the autocorrelations of simulated smFRET trajectories (**Figure S1**). With noise, however, only exponential fits the autocorrelations. Just as noise can convert a Lorentzian line shape into Gaussian line shape^53^, noise seems to turn a power law correlation into an exponential one. Note that the exponential fits recover the underlying sum of the simulated kinetic rates with and without noise.

### Larger inter-domain distance can accompany larger catalytic pocket opening

The catalytic cleft of MMP1 (∼0.5 nm) is narrow^17^, and as such, cleavage of substrates such as fibrin monomer (diameter ∼2-5 nm)^54^ and collagen monomer (diameter ∼1.5 nm)^55^ needs more opening of the catalytic pocket for easier access to the catalytic cleft. To test this, we used ANM simulations^56^ to calculate normal modes of MMP1 dynamics. ANM models proteins as a system of beads (amino acids) connected by springs (bonds) and calculates the preferred/normal modes of protein motion and corresponding eigenfrequencies of movement. Single molecule experiments (**Figure 3**) suggest that both active MMP1 and active site mutant of MMP1 has an open conformation (lower FRET) and a closed conformation (higher FRET). Therefore, we chose two MMP1 conformations (**Figure 5A** and **5B**) for ANM simulations (see Methods for details). To select the open conformation, we performed coarse-grained simulations of free MMP1 using ANM and decided on the MMP1 conformation with the highest inter-domain distance (**Figure 5A**). For closed conformation, we selected the conformation in the crystal structure (PDB ID: 1SU3) (**Figure 5B**). We considered 20 modes (20 frames for each mode) with the lowest frequencies (slower motion) to calculate the inter-domain distance (S142-S366) and catalytic pocket opening (N171-T230). A comparison of **Figure 5C** and **5D** shows that MMP1 in the open conformation has an overall catalytic pocket opening (∼2.7 nm), whereas the closed structure has an opening (∼2.6 nm). Since a larger catalytic pocket opening enables access to the catalytic cleft, we can infer that larger inter-domain distances (lower FRET) correlate with MMP1 activity.

**Figure 5.**
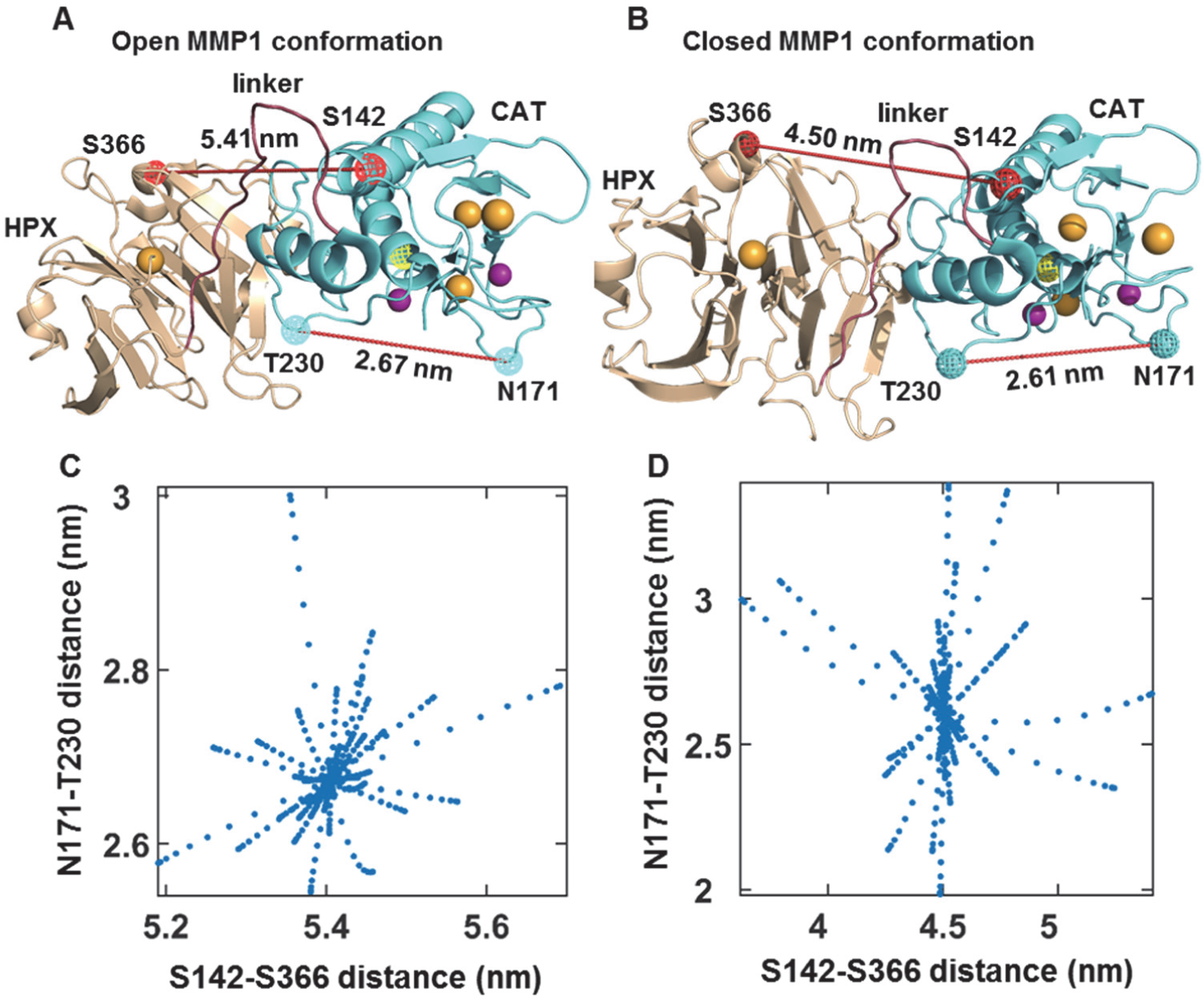
Correlations of MMP1 inter-domain distance with catalytic pocket opening when MMP1 is not bound to a substrate. Examples of (**A**) open and (**B**) closed conformations of MMP1. The inter-domain distances between S142 and S366 and corresponding catalytic pocketing openings between N171 and T230 have been noted. Spheres and cages represent the van der Waals radii. Yellow cage: E219; Wheat sphere: calcium; Mauve sphere: zinc. (**C**) and (**D**) are scatter plots of inter-domain distance and catalytic pocket opening for the open and closed conformations, respectively. Two distances are calculated using Anisotropic Network Modeling (ANM) simulations (see methods).

To investigate whether or not a larger inter-domain distance correlates with a larger catalytic pocket opening when MMP1 is bound to fibrin, we needed to determine the docking poses of MMP1 with fibrin because there is no crystal structure of fibrin-bound MMP1. To this end, open and closed conformations of MMP1 (**Figure 5A** and **5B**) were docked to a reconstructed model of fibrin using ClusPro^57^, a protein-protein docking server. The reconstructed model (see methods for detailed procedure) is a combination of the crystal structures of fibrinogen (PDB ID: 3GHG) and fibrin (PDB ID: 1FZC). The docking poses were sorted based on the distance from the α-carbon of E219 at the MMP1 catalytic site to the potential cleaving sites on fibrin. Since the cleavage sites on fibrin for MMP1 are unknown, we considered the cleavage sites for MMP3, MMP7, and MMP14 as substitutes: the α-chain at Asp97-Phe98, and Asn102-Asn103; the β-chain at Asp123-Leu124, Ans137-Val138, and Glu141-Tyr142; and the γ-chain at Thr83-Leu84^58^ (**Figure S2**). For each docking pose, we calculated the minimum distance possible for the three chains and considered the average of three lengths as the metric for the catalytic domain-fibrin proximity. In addition, we assumed that the hemopexin domain remains bound to the substrate, and thus, we considered the distance from the center of mass (CoM) of the hemopexin domain to any atom of the substrate as the metric for the hemopexin domain-fibrin proximity. We sorted the docking poses according to the two measures of proximity to the MMP1 domains. We selected the docking pose with minimum scores for the two criteria for ANM simulations (**Figure 6A** and **6B**). We considered 20 modes (20 frames for each mode) with the lowest frequencies (slower motion) to calculate inter-domain distance (S142-S366), catalytic pocket opening (N171-T230), and rms proximity between the MMP1 catalytic site and the three fibrin chains (**Figure 6C** and **6D**). The inter-domain distance reduces after binding fibrin for both the open and closed conformations of MMP1. As is the case of free MMP1 (**Figure 5C** and **5D**), fibrin-bound MMP1 also showed that larger inter-domain distance for the open conformation accompany larger catalytic pocket openings (**Figure 6C** and **6D**). In contrast, fibrinogen-bound MMP1 shows larger catalytic pocket opening for the closed conformation (**Figure S3**). The importance of open conformations on fibrin and closed conformations on fibrinogen is an experimentally testable insight from ANM simulations. At the single molecule level, we could test with fibrinogen attached to a quartz slide for TIRF imaging. Note that the overall charge is negative for both fibrinogen and a clean quartz slide, making it challenging to attach fibrinogen on a quartz slide. Nevertheless, histograms of smFRET values would be peaked at a higher FRET value on fibrinogen as compared to fibrin. One could also try to crystalize MMP1 with fibrinogen and fibrin, and check the conformations. Nevertheless, both fibrinogen- and fibrin-bound MMP1 show closer proximity to the three chains for the open conformation. The proximity suggests that a larger inter-domain distance is relevant for MMP1 activity.

**Figure 6.**
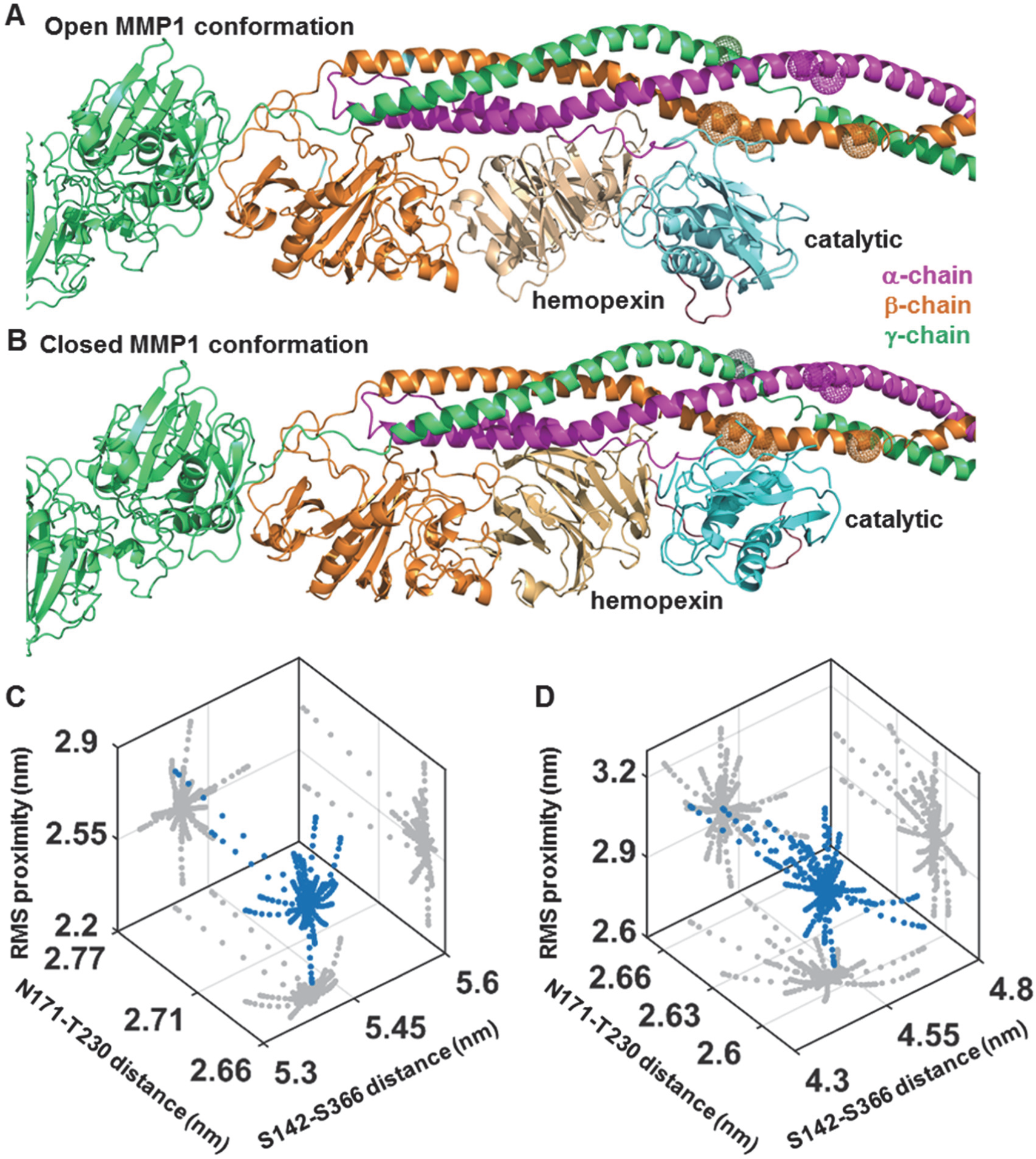
Correlations of MMP1 inter-domain distance with catalytic pocket opening when MMP1 is bound to the reconstructed fibrin model combining 3GHG and 1FZC. Examples of (**A**) open and (**B**) closed conformations of MMP1 bound to the reconstructed model of fibrin. Three dimensional scatter plots (blue circle) of inter-domain distance (S142-S366), catalytic pocket opening (N171-T230), and rms proximity between the MMP1 catalytic site and the three fibrin chains for (**C**) open and (**D**) closed MMP1 conformations. Two-dimensional projections of the scatter plots are in gray. The open structure shows larger catalytic pocket openings.

### The proximity of MMP1 to the three chains correlates with the sequence of degradation

Since MMP1 needs to come in the proximity before cleavage, we performed molecular docking of MMP1 with fibrin. **Figure 7A** shows MMP1 molecular docking poses superimposed on the reconstructed model of fibrin. MMP1 binds to fibrin at specific places in both open and closed conformations. To gain further insights, we calculated the distance between every atom in MMP1 and the three fibrin chains. We counted the number of distances below 0.5 nm for the top 30 docking poses and calculated the mean and standard deviation (**Figure 7B**). On average, MMP1 in open conformation showed the closest proximity to the α-chain (**Figure 7B**, left panel). We quantified the statistical significance by p-value (**Figure 7B**, left panel). In contrast, the closed conformation of MMP1 did not show any significant difference in proximity to the three chains (**Figure 7B**, right panel).

**Figure 7.**
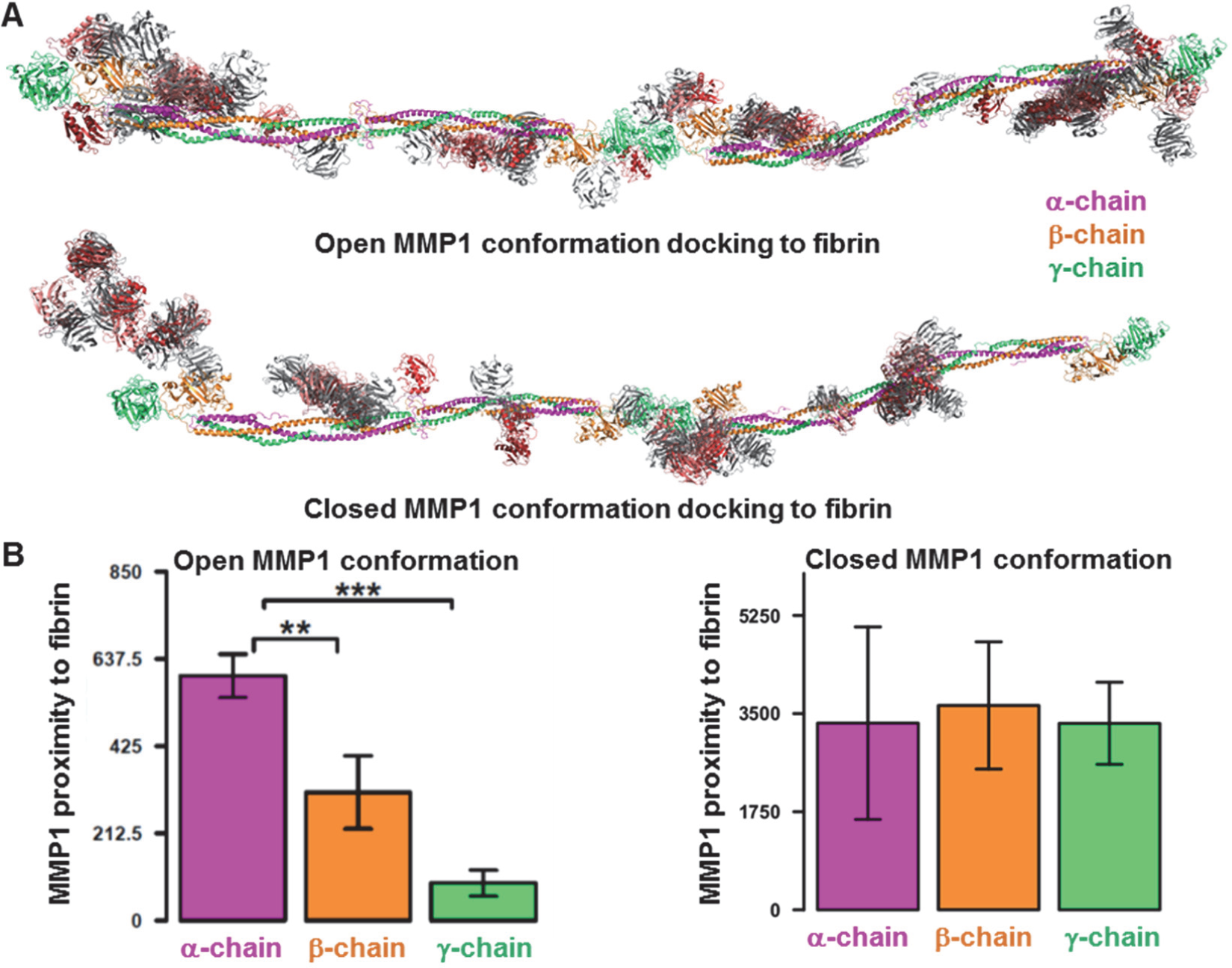
Proximity of MMP1 to the chains of reconstructed fibrin model. (**A**) The clustered 30 docking poses obtained from ClusPro using fibrin as the ligand and MMP1 as the receptor. The catalytic domain of MMP1 is red, whereas the hemopexin domain is gray. (**B**) We measured the distance between all possible pairs of atoms between the fibrin chains and MMP1 for 30 docking poses obtained from ClusPro. We counted any distance less than 5 Å and plotted the distributions of total count (y-axis) for each chain for the open and closed conformations. p-value<0.05:*; p-value<0.01:**; p-value<0.001:***.

To test whether or not the sequence of MMP1 proximity to the three chains has any implication in fibrinolytic activity, we performed the time-dependent degradation of fibrinogen. **Figure S4B** shows that the overall catalytic activity is fastest on the α-chain and slowest on the γ-chain. Fibrinogen, the basic building block of fibrin, has three pairs of amino acid chains: 1) the α-chain at ∼63 kDa, 2) the β-chain at ∼56 kDa, and 3) the γ-chain at ∼47 kDa. SDS PAGE of fibrinogen control approximately confirms molecular weights of the three chains (**Figure S4A**, lane 2 from left). These three chains are connected by a dimeric disulfide knot (DSK) at the N-termini and many other disulfide bonds along the length of fibrinogen^59,60^. Treatment of 1 mg/mL of fibrinogen with 0.1 mg/mL of active MMP1 at 37° C for 24 h resulted in cleavage of all three chains and a final prominent fragment at ∼40 kDa (**Figure S4A**, lane 3 from left). MMP1 completely degraded the α-chain and partially degraded the β-chain within 2 h; however, the γ-chain took nearly 6 h to be degraded (**Figure S4B**). After 8 h, a prominent band at ∼40 kDa remained. In other words, the computationally determined sequence of degradation in the open conformation agrees with the experimentally observed sequence (**Figure S4B**). The agreement between the computational and experimental sequence of degradation further confirms that the open conformations observed in smFRET experiments are functionally relevant.

We also imaged fibrin without crosslinks after treatment with MMP1 using SEM (**Figure S4C**), which shows increased porosity of fibrin for active MMP1 consistent with the degradation. We used fibrin without crosslinks because it is easier to form a thin layer of fibrin on a quartz slide for smFRET measurements, and the crystal structure without crosslinks (PDB ID: 1FZC) is available. However, fibrin with crosslinks is more physiologically relevant (see methods for making fibrin with crosslinks). As such, we quantified the degradation rate of fibrin with crosslinks using a weight-based assay (Figure S5). We also performed gel electrophoresis of degradation product (**Figure S6A**), imaged the reaction volume to show the degradation within 5 h (**Figure S6B**), and imaged the surface morphology using SEM (**Figure S6C**).

### Unlike collagen, MMP1 domains do not communicate via fibrin

Previously, we performed ANM simulations of MMP1 interacting with collagen using the crystal structure of MMP1 bound to a collagen model (4AUO)^23^. We reported that the catalytic and hemopexin domains could communicate via collagen even if the physical linker is removed^23^, which explained the experimental observation that a mixture of the two domains purified separately could degrade triple-helical collagen^18^. We wanted to investigate whether or not the two domains communicate via fibrin. We created three forms of MMP1 using the crystal structure (PDB ID: 1SU3): 1) full-length MMP1, 2) MMP1 with the linker domain removed, and 3) MMP1 with the linker and hemopexin domain removed. We docked both the open and closed conformations with the reconstructed fibrin using ClusPro and chose the binding pose shown in **Figure 8**. Three forms of MMP1 interacting with fibrin resulted in similar mean and standard deviation of the catalytic pocket opening (**Figure 8**). In contrast, the catalytic pocket opening on collagen changed even without the physical linker^23^. In other words, fibrin interacts with MMP1 passively, whereas collagen interacts with MMP1 actively. Using single molecule tracking of labeled MMP1 on type-1 collagen fibrils, we also showed that fibrils play a role in MMP1 activity due to vulnerable sites on fibrils created during fibril assembly^61^.

**Figure 8.**
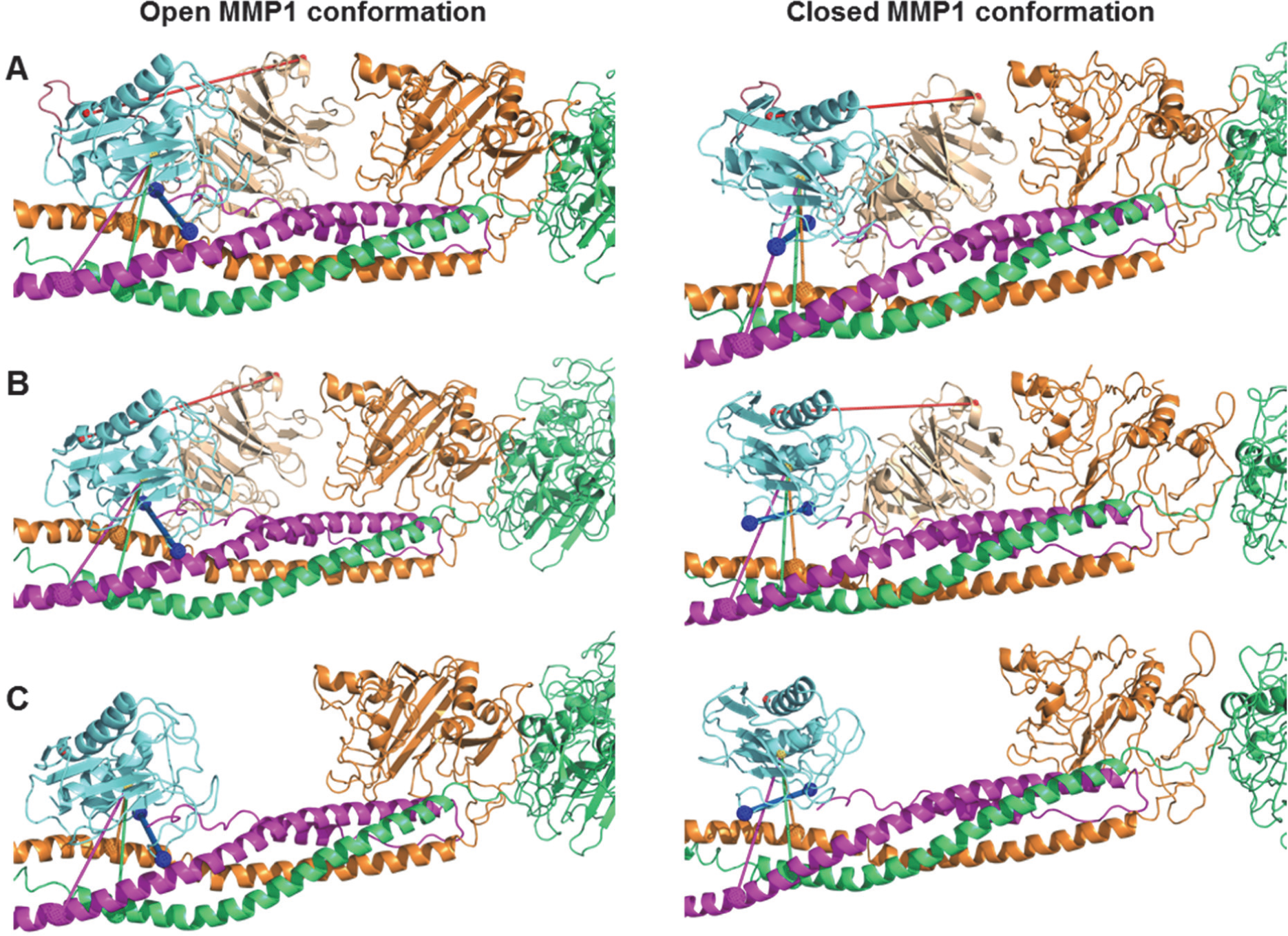
Effects of the linker and reconstructed fibrin on the catalytic pocket opening. The catalytic pocket openings on fibrin represented by the blue line connecting N171 and T230 for (**A**) the full-length MMP1 (open conformation: 2.68±0.01 nm; closed conformation: 2.61±0.01 nm); (**B**) the two MMP1 domains without the linker (open conformation: 2.68±0.01 nm; closed conformation: 2.61±0.01 nm); and (**C**) the catalytic domain alone (open conformation: 2.68±0.01 nm; closed conformation: 2.68±0.01 nm). The error bars represent the standard deviation of 60 measurements of the catalytic pocket opening obtained from 20 frames each for the three slowest normal modes.

Both fibrin and collagen are extracellular and triple-helical but have microstructural and mechanical differences^62^. Fibrin monomers are 45 nm long and have three right-handed chains forming a left-handed structure in between two globular regions^54^. In contrast, collagen monomers are 300 nm long and have three left-handed chains forming a right-handed structure^63^. The difference in handedness between fibrin and collagen may lead to in-phase and out-of-phase motion of MMP1 domains. To understand why fibrin and collagen interact differently, however, we need further studies since fibrin and collagen have different sequences and structures despite the triple-helical similarities.

In summary, we have shown that the inter-domain dynamics of MMP1 on fibrin correlates with its fibrinolytic activity. We measured smFRET between two fluorophores attached to the catalytic and hemopexin domains of MMP1. A comparison of distributions of smFRET values between active MMP1 and active site mutant of MMP1 suggests that MMP1 needs to have the two domains well-separated for function. To investigate the roles of open conformations, we measured smFRET in the presence of MMP9 and tetracycline. MMP9 is an enhancer, whereas tetracycline is an inhibitor of MMP1 activity on collagen and a synthetic substrate. In the presence of tetracycline, the inter-domain distance on fibrin becomes smaller consistent with the trend on collagen. MMP9 causes the inter-domain distance of MMP1 to be shorter on fibrin, which is opposite to the observation on collagen. One possible reason is that MMP9 by itself can degrade fibrin but cannot degrade triple-helical collagen. As a result, fibrin degradation by MMP1 in the presence of MMP9 is more complicated. A two-state Poisson process quantitatively describes the inter-domain dynamics. We fitted the histograms of smFRET values to a sum of two Gaussians. The best-fit parameters for the centers are the two states used for simulations. The ratio of kinetic rates equals to the ratio of areas of the two Gaussians fitted to a histogram. Additionally, the sum of kinetic rates equals to the decay rate of autocorrelations of smFRET values. We simulated smFRET trajectories with experimentally determined parameters and noise levels. We recovered the underlying parameters to validate our analyses. The presence of noise in the signal appears to convert a power law autocorrelations into an exponential one.

We performed molecular docking of the open and closed conformations of MMP1 with fibrin. In the closed conformation, MMP1 approaches the three chains equally. In the open conformation, however, MMP1 approaches the α-chain first and the γ-chain last, which we confirmed using SDS PAGE of fibrinogen degraded by MMP1. As such, the open conformation appears to be more functionally relevant. One or both domains of MMP1 binds to fibrin at many places along the length of fibrin. However, the primary location on fibrin is near the globular regions where both the domains bind. We, therefore, selected the docking pose near the globular part for ANM simulations. ANM simulations of MMP1-fibrin interactions revealed that the open conformation with well-separated domains has a larger catalytic pocket opening. A larger opening enables the fibrin chains to approach closer to the MMP1 active site. In the closed conformation, the fibrin chains get closer to the catalytic site only when the catalytic domain is alone. We performed limited digestion of fibrin by MMP1 with and without crosslinks and observed increased porosity due to degradation. Also, MMP1 degrades and dissolves crosslinked fibrin within 5 hr, suggesting a role of MMP1 in the fibrinolytic pathway. We quantified the degradation using a weight-based assay because interactions of water-soluble MMP1 with water-insoluble fibrin are challenging to quantify using solution-based biochemical assays.

We discussed the results of the MMP1-fibrin system in light of our previous studies on the MMP1-collagen system. For both fibrin and collagen, the open conformation of MMP1 is functionally relevant and responds to tetracycline similarly. The open conformation enables larger catalytic pocket opening and closer proximity to the three strands of fibrin and collagen. A two-state Poisson process describes MMP1 dynamics on both fibrin and collagen. However, MMP1 dynamics are opposite to each other on fibrin and collagen in the presence of MMP9. Note that MMP9 can degrade fibrin but cannot degrade triple-helical collagen and, as such, can influence MMP1 dynamics differently. Collagen plays an active role in mediating allosteric communications between the two domains and opening the catalytic pocket. In contrast, fibrin plays a passive role and cannot mediate communications between the two domains. Further studies are needed to investigate the detailed mechanisms of different MMP1 interactions on fibrin and collagen, which may enable allosteric control of MMP1 activity in a substrate-dependent manner for exploring MMPs as drug targets.

## Methods

### Purification and labeling of MMP1

Full-length recombinant MMPs were expressed in *E. coli* and activated using 0.1 mg/mL trypsin. Activated MMP1 and trypsin together create a chain reaction of further MMP1 activation and broad-spectrum proteolytic cleavage of *E. coli* proteins. Molecular weights of activated MMP1 and trypsin are ∼43 kDa and ∼23 kDa, respectively. We used a 30 kDa cut-off Amicon filter to centrifuge and purify MMP1 in the activated form. For further details, see the previous publication on protease-based purification method^14^. We labeled purified MMP1 with AlexaFluor555 (ThermoFisher Scientific, Cat# A20346, donor dye) and AlexaFluor647 (ThermoFisher Scientific, Cat# A20347, acceptor dye) in a ratio of ∼1:1 using maleimide chemistry. 1 mL of MMP1 at 1 mg/mL concentration was incubated with 20 µL of 1 mg/mL AlexaFluor555 and AlexaFluor647 for 1 h in a 5 mL glass vial in a nitrogen environment to avoid oxidation of the dyes. After incubation, we used a 30 kDa cut-off Amicon filter to remove free dyes from the solution. We compared the specific activities of labeled and unlabeled MMP1 on the synthetic substrate, MCA-Lys-Pro-Leu-Gly-Leu-DPA-Ala-Arg-NH2, (R&D Systems, Cat# ES010) as described before^14^. Labeling MMP1 does not affect its specific activity^23^.

### Single-molecule measurements

We prepared 200 µL of reaction volume in 10 mM Phosphate Buffer Saline (PBS) (Sigma, Cat# P3813, pH 7.4) by mixing 10 units of thrombin (Cayman chemical, Cat# 13188) and 50 μg of fibrinogen (Cayman chemical, Cat# 16088). A thin layer of fibrin was created on a quartz slide similar to the blood smear protocol used in diagnostics^64^ and incubated at 37° C for 18 h without shaking. We made a flow cell of thickness 120 μm using a piece of double-sided adhesive tape, a clean quartz slide, and a glass coverslip. The quartz slides had two holes to connect the input and output tubes for exchanging buffers and solutions. We created an evanescent wave at the interface of the quartz slide and sample in a Total Internal Reflection Fluorescence (TIRF) Microscope as described before^65-68^. We incubated 50 µL of 0.1 mg/mL labeled MMP1 with 50 µL of protein buffer (50 mM Tris, 100 mM NaCl, pH 8.0) and 100 µg/mL tetracycline for 30 min at 22° C. The labeled MMP1 was serially diluted to prepare a working concentration of ∼100 pM and flowed into the flow cell. Alexa555 dyes (donor dyes) attached to MMP1 was excited using a 532 nm laser and imaged using the TIRF microscope at 22° C at 100 ms time resolution. We imaged Alexa647 and Alexa555 emissions using an EMCCD camera (Andor iXon). The two emission channels were superimposed using a pairwise stitching plugin of ImageJ, and the intensities from the two dyes were extracted and analyzed using Matlab.

### Reconstruction of the extended fibrin molecule

We superimposed the crystal structure of fibrinogen (PDB ID: 3GHG)^69^ with the crystal structure of D-D fibrin fragment (PDB ID: 1FZC)^70^ using the Combinatorial Extension (CE) algorithm^71^ implemented in PyMOL^72^ to obtain the reconstructed model of fibrin. The fibrinogen asymmetric unit presents two replicas (either the chains A-F+M-P or G-L+Q-T). Each replica has two ends, which are represented by chains A-C and chains D-F in one and by chains G-I and J-L in the other. Each of such fibrinogen ends could be superimposed to any of the ends of the fibrin crystal (either the chains A-C or chains D-F), yielding a total of 16 different possibilities to reconstruct the fibrin molecule. For each of the superimposed assembly, we superimposed a new fibrin crystal to the fibrinogen biological assembly not used for the initial superimposition. Thus, we superimposed two extra fibrin crystals (Fa and Fb, one per superimposed fibrinogen biological assembly) on each of the 16 different reconstruction attempts. A reconstruction compatible with the original fibrinogen packing would show no differences in the positioning of Fa and Fb. Thus, to select the best superimposition combination to reconstruct the fibrin molecule, the one that minimized the RMSD in between Fa and Fb was selected.

### Selection of closed and open conformations of MMP1

The activation peptide (residues 32 to 98) was removed from the first biological assembly (chain A) of the crystal structure of MMP1 (PDB ID: 1SU3), leaving only the catalytic (residue numbers 107 to 260), linker (261-277) and hemopexin (278-466) domains of the enzyme, which we selected as the closed conformation. To obtain the open conformation, we submitted the coordinates of the closed conformation to the ANM web server 2.1^56^ using default parameters. We extracted the coordinates of the three-dimensional fluctuations for 20 frames of the 20 slowest modes. For each frame on each model, we calculated distances using PyMOL. We used the distance between S142 and S366 to analyze the fluctuations between the catalytic and hemopexin domains. We selected the frame that maximized this distance (second mode, second frame) as the MMP1 open conformation.

### Molecular docking of MMP1 with fibrin

The closed and open conformations of MMP1 were docked individually with the reconstructed fibrin molecule using the default parameters of ClusPro 2.0 web server^57^. We used the reconstructed fibrin as the receptor and the MMP1 conformations as the ligands. We considered the centroid structure of each of the resulting clusters (maximum 30) as the representative docking solution for a cluster, and we obtained the corresponding weighted scores for the best coefficient for each binding pose. We adjusted different poses to the same Cartesian space by aligning the reconstructed fibrin from each pose to a template one using PyMOL. To analyze the accessibility of MMP1 to the three chains, we counted the number of times an atom in MMP1 was within 5 Angstrom from an atom in the chains.

### Preparation of fibrin without and with crosslinks

We prepared fibrin without crosslinks by adding 10 units of thrombin (Cayman chemical, Cat# 13188) to 50 μg of fibrinogen (Cayman chemical, Cat# 16088) in 10 mM PBS to a total reaction volume of 200 μL and incubating at 37° C for 18 h with 250 rpm shaking inside an incubator (Thermo Fisher Scientific, MaxQ600). Preparation of crosslinked fibrin involved two additional reagents: factor XIII (Abcam, Cat # ab62427) and CaCl_2_. Thrombin activates factor XIII to factor XIIIa, which in the presence of Ca^++^ induces crosslinks in fibrin. We added 10 units of thrombin to 2.5 µg of factor XIII in a 0.5 mL PCR tube, diluted with 10 mM PBS to a total reaction volume of 100 μL, and incubated at 37° C for 10 min. After incubation, we added 50 μg of fibrinogen and 10 µL of 5 mM CaCl_2_, diluted with PBS buffer to a total reaction volume of 200 µL, and incubated at 37° C for 15 min to obtain gel-like crosslinked fibrin. Note that fibrin without crosslinks requires a longer incubation time and shaking, but fibrin with crosslinks forms within 15 min without shaking. For every reaction, we put a part of the sample on a sample mount before drying and imaging the surface morphology of fibrin using a Phenom Pro Scanning Electron microscope.

### Sodium Dodecyl Sulphate-Polyacrylamide Gel Electrophoresis (SDS PAGE)

The degradation profile after fibrinolysis was analyzed using SDS-PAGE. We mixed SDS PAGE loading buffer containing BME (Biorad, Cat# 610737) with each sample in a 1:1 (v/v) ratio and incubated at 95° C for 10 min. We loaded the samples onto the wells of 8% Tris.glycine gels. We ran electrophoresis in Tris.glycine buffer (Biorad, Cat# 1610732) for 30 min at 50 V, followed by 1 h at 120V. We used the precision plus protein kaleidoscope pre-stained protein standards (Biorad, Cat# 1610375) as molecular weight markers. The SDS-PAGE gel was stained with Coomassie Brilliant Blue R-250 (Biorad, Cat# 161-0436) and imaged using a digital camera after destaining.

## Acknowledgments

One grant to S.K.S. and J.K.S. from the National Institutes of Health (RGM137295A) partially supported this work. This project was also partially funded by the professional development funds given to S.K.S and J.K.S. at the Colorado School of Mines. JPI was supported by the grant MSCAfellow@MUNI.

## Author contributions

S.K.S. conceived and designed the overall project. L.K. and S.K.S. designed experiments, L.K. performed experiments, J.P. performed docking and ANM analysis. L.K., J.P., J.K., C.H., J.K., D.W., and S.K.S. analyzed data. S.K.S. wrote the manuscript. All authors edited the manuscript.

## Data availability

The datasets generated during and/or analyzed during the current study are available from the corresponding author on reasonable request.

## Supplementary Information

We fitted a sum of two Gaussians to the experimental histograms using the following equation:

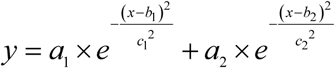

where a, b, and c are amplitude, center, and width of the Gaussian, respectively. The parameters b1 and b2 are the two states, S1 and S2, respectively.

We subtracted the average FRET value from each FRET trajectory and used the following equation to calculate autocorrelations:

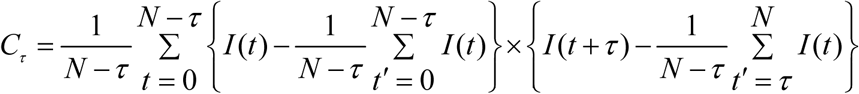

where *C*_*τ*_ is the autocorrelation at lag number *τ, N* is the number of points in a FRET trajectory, and *I* (*t*) is the FRET value at *t*.

**Table S1.** Best-fit parameters for experimental histograms and autocorrelations. (**A**) A sum of two Gaussians fits the experimental histograms in **Figure 3**. (**B**) An exponential distribution fits the experimental autocorrelations in **Figure 3**. Power law distribution does not fit the experimental autocorrelations. (**C**) Kinetic rates of interconversion between the two states, S1 and S2, from the histograms and autocorrelations. Yellow highlighted numbers are (A) the centers of Gaussian fits and (B) the decay rates of exponential fits, e. The error bars represent the standard errors of mean.

A Gaussian fit parameters for experimental histograms
| MMP1 without ligands |  |  | MMP1 with MMP9 |  | MMP1 with tetracycline |  |
| --- | --- | --- | --- | --- | --- | --- |
|  | Active | Inactive | Active | Inactive | Active | Inactive |
| a1 | 1.99±0.13 | 0.90±0.17 | 2.14±0.06 | 1.18±3.21 | 4.38±0.05 | 0.60±0.79 |
| b1/S1 | 0.42±0.01 | 0.52±0.01 | 0.46±0.01 | 0.46±0.18 | 0.61±0.01 | 0.49±0.13 |
| c1 | 0.08±0.01 | 0.10±0.01 | 0.11±0.01 | 0.11±0.06 | 0.10±0.01 | 0.11±0.06 |
| a2 | 5.52±0.15 | 8.12±0.17 | 5.58±0.07 | 4.61±4.12 | 2.10±0.12 | 5.6±1.21 |
| b2/S2 | 0.51±0.01 | 0.55±0.01 | 0.51±0.01 | 0.53±0.01 | 0.70±0.01 | 0.58±0.01 |
| c2 | 0.07±0.01 | 0.06±0.01 | 0.06±0.01 | 0.09±0.01 | 0.06±0.01 | 0.09±0.01 |

B Exponential fit parameters for correlations
| MMP1 without ligands |  |  | MMP1 with MMP9 |  | MMP1 with tetracycline |  |
| --- | --- | --- | --- | --- | --- | --- |
|  | Active | Inactive | Active | Inactive | Active | Inactive |
| d | 0.37±0.01 | 0.15±0.01 | 0.21±0.01 | 0.45±0.01 | 0.62±0.01 | 0.36±0.01 |
| e | 0.08±0.01 | 0.07±0.01 | 0.09±0.01 | 0.05±0.01 | 0.05±0.01 | 0.06±0.01 |
| f | -0.03±0.01 | -0.01±0.01 | -0.01±0.01 | -0.03±0.01 | -0.05±0.01 | -0.004±0.001 |

C Kinetic rates calculated from histograms and correlations
| MMP1 without ligands |  |  | MMP1 with MMP9 |  | MMP1 with tetracycline |  |
| --- | --- | --- | --- | --- | --- | --- |
|  | Active | Inactive | Active | Inactive | Active | Inactive |
| k1 (s<sup>-1</sup>) | 0.0567 | 0.0591 | 0.0528 | 0.0381 | 0.0112 | 0.0531 |
| k2 (s<sup>-1</sup>) | 0.0233 | 0.0109 | 0.0372 | 0.0119 | 0.0388 | 0.0069 |

We normalized autocorrelations by dividing autocorrelations at each lag by *C*_*τ* =0_. We fitted autocorrelations between *τ* = 1 and *τ* = 1000 to both power law and exponential distributions. For power law, we used a form of Pareto distribution^1^ that satisfies the boundary conditions of our calculated autocorrelations, i.e., *C*_*τ* = 0_ = 1at *t* = 0 and *C*_*τ* = ∞_ = 0 at *t* = ∞. We fitted the following equations of power law and exponential functions:

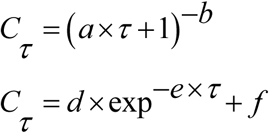

### Best-fit parameters for two-state simulations

We simulated smFRET trajectories assuming that MMP1 undergoes interconversion between two states, S1 and S2, which are the centers of Gaussian fits to the histograms. We considered active MMP1 and active site mutant of MMP1 without ligands (**Figure 3A**). We simulated 350 smFRET trajectories, each 1000 s long, with the input parameters in **Table S2**. We analyzed the simulated and experimental trajectories similarly. The recovered parameters (**Table S2**, right side) agree well with the input parameters.

**Table S2.**
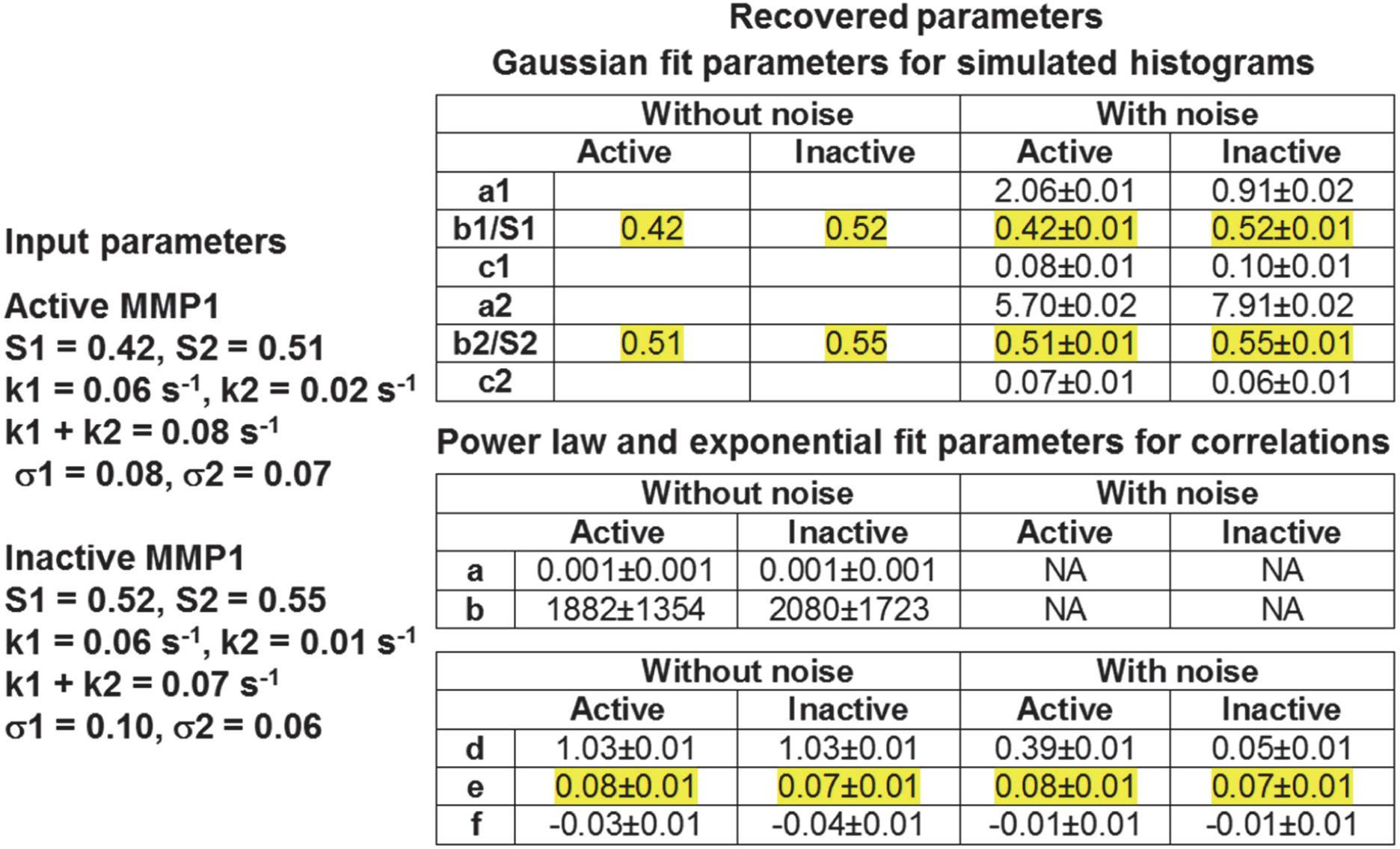
MMP1 inter-domain dynamics as a two-state Poisson process. Analysis of FRET trajectories for MMP1 without ligands recovered the centers and decay rates used as inputs. The agreement between the input and retrieved values suggests that a two-state Poisson process describes the MMP1 dynamics.

**Figure S1.**
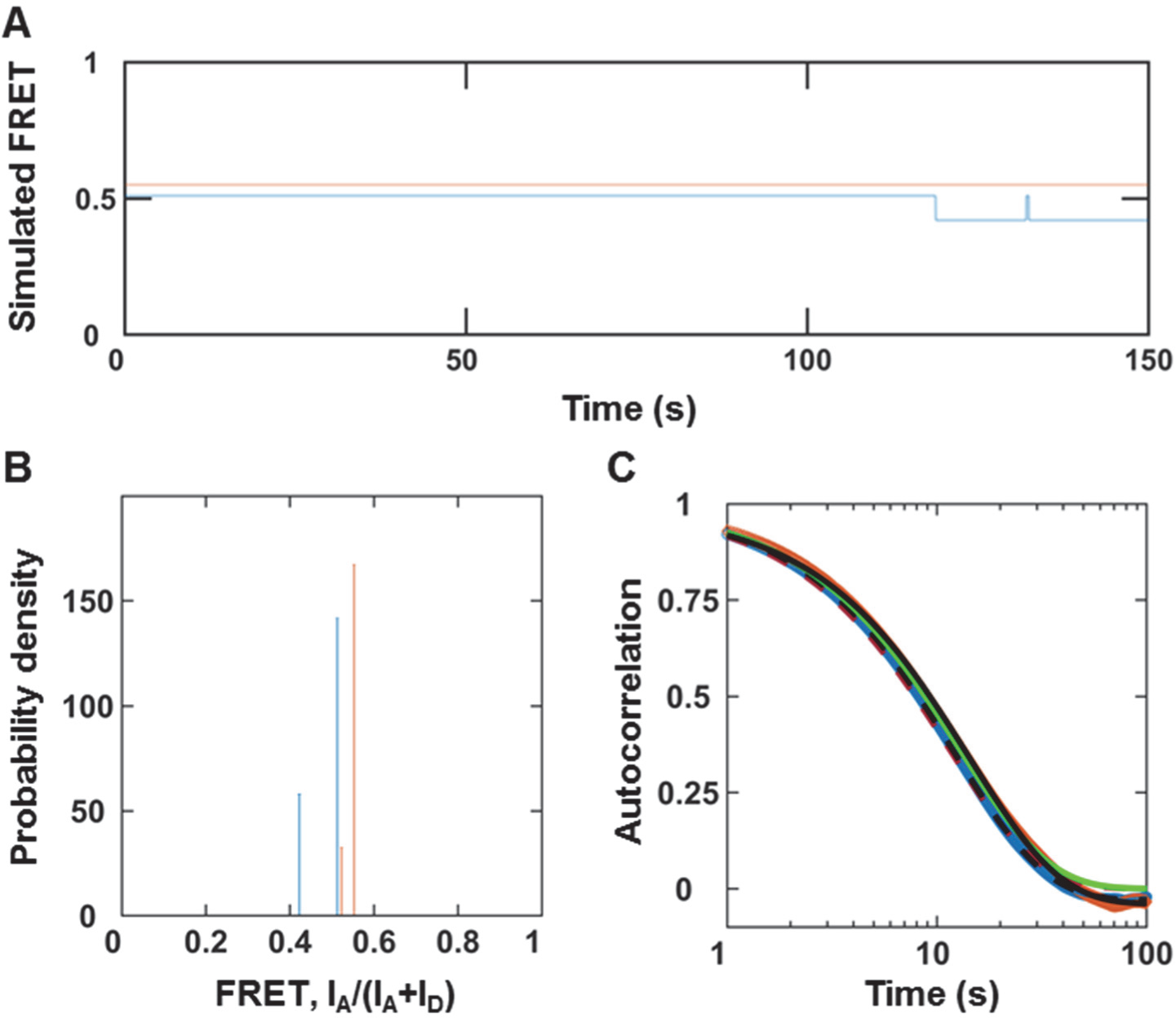
MMP1 inter-domain dynamics as a Poisson process without noise. (**A**) An example of simulated two-state FRET trajectory without noise for active MMP1 (blue) and active site mutant of MMP1 (orange). (**B**) Histograms of the recovered FRET values with bin size=0.005. (**C**) Autocorrelations of simulated trajectories recover the sum, k1+k2, from exponential fits (active MMP1: dashed black line; active site mutant of MMP1: solid black line). Note that power law does not fit the autocorrelations with noise (**Fig. 4**). However, power law fits the autocorrelations without noise (active MMP1: dashed red line; active site mutant of MMP1: solid green line).The error bars are the sems for histograms and autocorrelations and are too small to be seen.

### Potential sites of interactions between MMP1 and fibrin

MMP3, MMP7, and MMP14 cleave the α-chain at Asp97-Phe98 and Asn102-Asn103; the β-chain at Asp123-Leu124, Asn137-Val138, and Glu141-Tyr142; and the γ-chain at Thr83-Leu84^2^ as shown in **Figure S2**.

**Figure S2.**
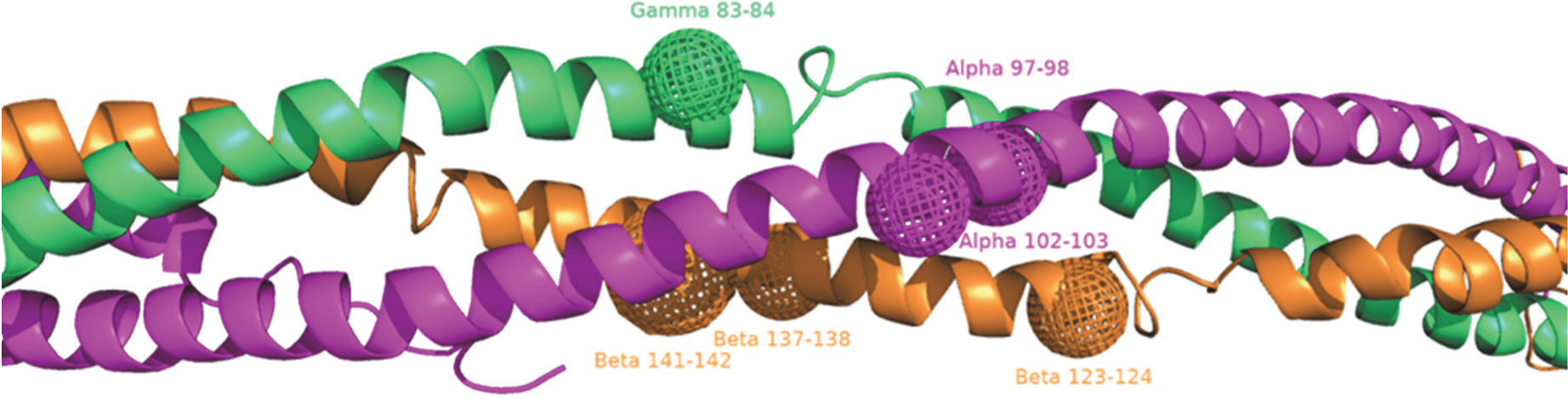
Potential MMP1 cleavage sites on the three fibrin chains. The balls are potential cleavage sites on fibrin that we considered to select the best docking pose between MMP1 and fibrin.

### Inter-domain distance correlates with the catalytic pocket opening of MMP1 on fibrinogen

In contrast to free MMP1 (**Figure 5**) and fibrin-bound MMP1 (**Figure 6**), fibrinogen-bound MMP1 (**Figure S3**) shows smaller catalytic pocket openings for the open MMP1 conformations. However, fibrinogen-bound MMP1 shows closer proximity to the fibrinogen chains in agreement with free MMP1 and fibrin-bound MMP1. The smaller catalytic pocket opening suggests a different mechanism of fibrinogen and fibrin degradation by MMP1, which needs further studies to confirm.

**Figure S3.**
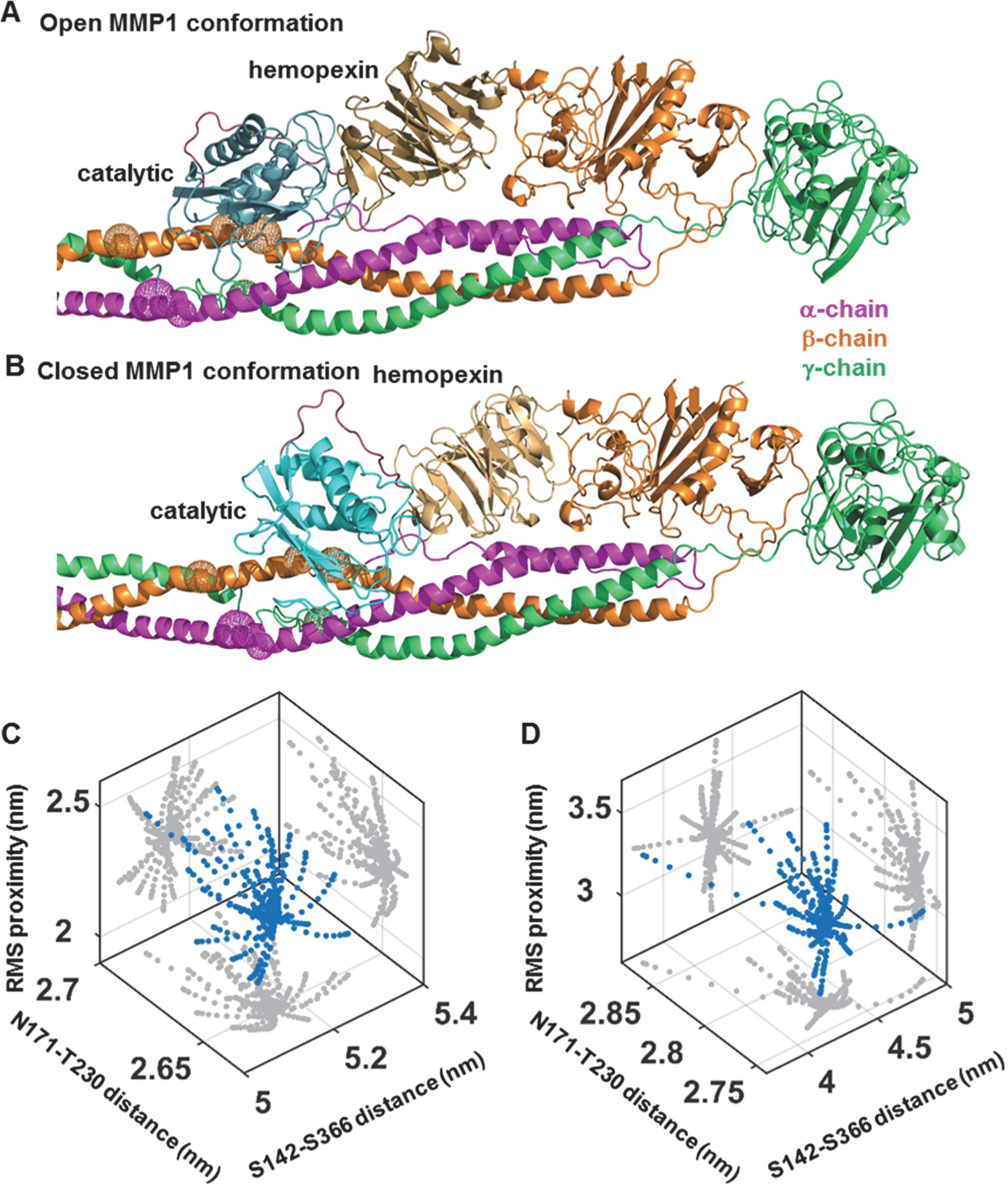
Correlations of MMP1 inter-domain distance with catalytic pocket opening when MMP1 is bound to fibrinogen. Examples of (**A**) open and (**B**) closed conformations of MMP1 (PDB ID: 1SU3) bound to fibrinogen (PDB ID: 3GHG). Three dimensional scatter plots (blue circle) of inter-domain distance (S142-S366), catalytic pocket opening (N171-T230), and rms proximity between the MMP1 catalytic site and the three fibrinogen chains for (**C**) open and (**D**) closed MMP1 conformations. Two-dimensional projections of the scatter plots are in gray.

### MMP1 activity on fibrinogen

The point mutation E219Q renders MMP1 catalytically inactive on collagen. For fibrinogen degradation, however, the E219Q mutant degraded the α- and β-chains, but did not degrade the γ-chain (**Figure S4A**, lane 4 from left). We used trypsin during MMP1 purification, and trypsin is known to degrade fibrinogen^3^; therefore, any residual trypsin could potentially interfere with the results. To address this possibility, we added 0.5 mg/mL trypsin inhibitor to all the reactions, which was sufficient to inhibit 0.1 mg/mL trypsin (**Figure S4A**, lane 5 from left).

**Figure S4.**
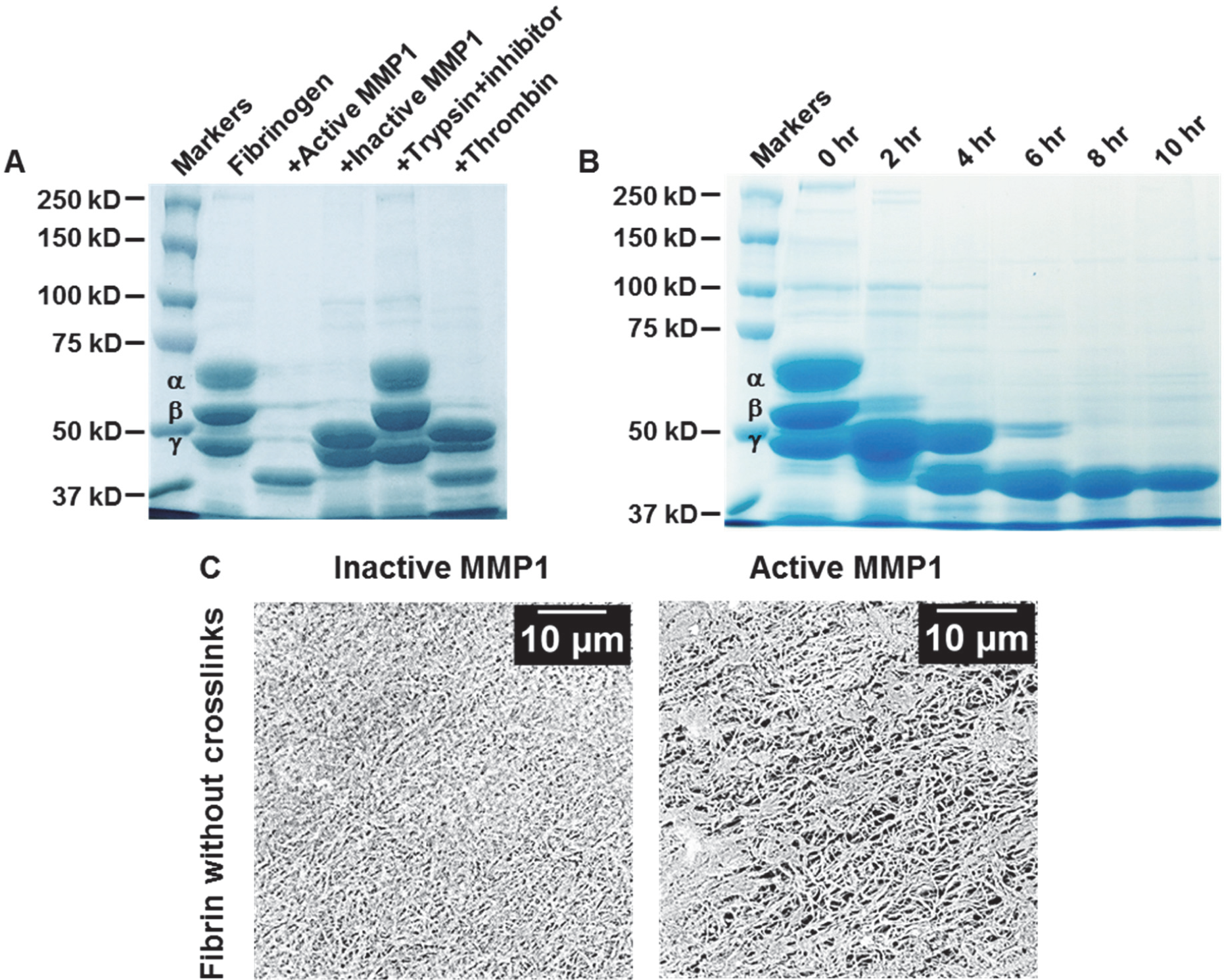
Ensemble activity of MMP1 on fibrinogen and fibrin without crosslinks. (**A**) SDS PAGE of fibrinogen treated with different enzymes. Since we used trypsin to activate and purify MMP1, we added 0.5 mg/mL trypsin inhibitor to lane 2-6 to control for any residual effect of trypsin. Trypsin inhibitor (Worthington, Lima Bean, Cat# LS002829) at 0.5 mg/mL inhibited the fibrinolytic effect of 0.1 mg/mL trypsin used in lane 5. Note that the difference between trypsin inhibitor and MMP1 inhibitor (tetracycline). (**B**) Time-dependent fibrinogen degradation by active MMP1. (**C**) SEM images of fibrin after MMP1 treatment. Note that the active site mutant of MMP1 is catalytically inactive on collagen^4^. However, as evident from **Figure S4A**, active site mutant of MMP1 is partially active on fibrinogen. In contrast, active site mutant of MMP1 is catalytically inactive on crosslinked fibrin, as shown in **Figure S5** and **Figure S6**.

Even though limited digestion of fibrinogen by thrombin leads to fibrin, the longer digestion for 24 h led to further degradation of fibrinogen (**Figure S4A**, lane 6 from left). **Figure S4A** does not reveal the sequence in which MMP1 degrades the three chains of fibrinogen. **Figure S4B** shows the time-dependent degradation of fibrinogen. MMP1 degrades the α-chain first and the γ-chain last. A comparison with prior research shows that plasmin, the well-known fibrinolytic agent, also degrades the α-chain first and the γ-chain last^5^. Next, we tested the effects of MMP1 on fibrin morphology using SEM (**Figure S4C**), where the destruction of fibrin structures is visible for active MMP1. We created similar thin layers of fibrin morphologies for smFRET experiments. Overall, the ensemble experiments in **Figure S4** show that MMP1 has fibrinolytic activity on fibrinogen and fibrin, and there is a specific sequence of degradation of the three chains. Both thrombin and trypsin cleave the same Arg-Gly bonds in fibrinogen^6^. However, thrombin is highly specific, and only four bonds linking the fibrinopeptides to the rest of the fibrinogen molecule are rapidly hydrolyzed by thrombin, whereas trypsin cleaves all peptide bonds in fibrinogen that follow arginine or lysine^6^.

In comparison, degradation of fibrinogen and crosslinked fibrin by MMPs do not have specific cleavage sites across the MMP-family. For example, MMP3, MMP7, and MMP14 cleave the α-chain at Asp97-Phe98 and Asn102-Asn103; the β-chain at Asp123-Leu124, Asn137-Val138, and Glu141-Tyr142; and the γ-chain at Thr83-Leu84^2^. Cleavage sites on both fibrinogen and crosslinked fibrin for these three MMPs are close to those for plasmin. Additionally, MMP3 solubilizes crosslinked fibrin by cleaving the Gly404-Ala405 bond in the γ-chain^7^, resulting in a D-like monomer fragment at ∼94 kDa similar to fibrinogen degradation by plasmin. In contrast, MMP7 and MMP14 solubilize crosslinked fibrin and produces D-like dimer fragments at ∼186 kDa similar to crosslinked fibrin degradation by plasmin^2^. However, MMP1, MMP2, MMP9, and MMP15 do not show similar activity^2^. Fibrinogen degradation by catalytic domains of MMP8, MMP12, MMP13, and MMP14 show differences in the cleavage sites^8^. Overall, MMPs show differences in fibrinolytic activity, which is intriguing because the catalytic domain has a largely conserved sequence among the MMP-family members^9^. Interestingly, prior research reported insignificant fibrinolytic activity of MMP1^7^, in contrast to our findings.

**Figure S5.**
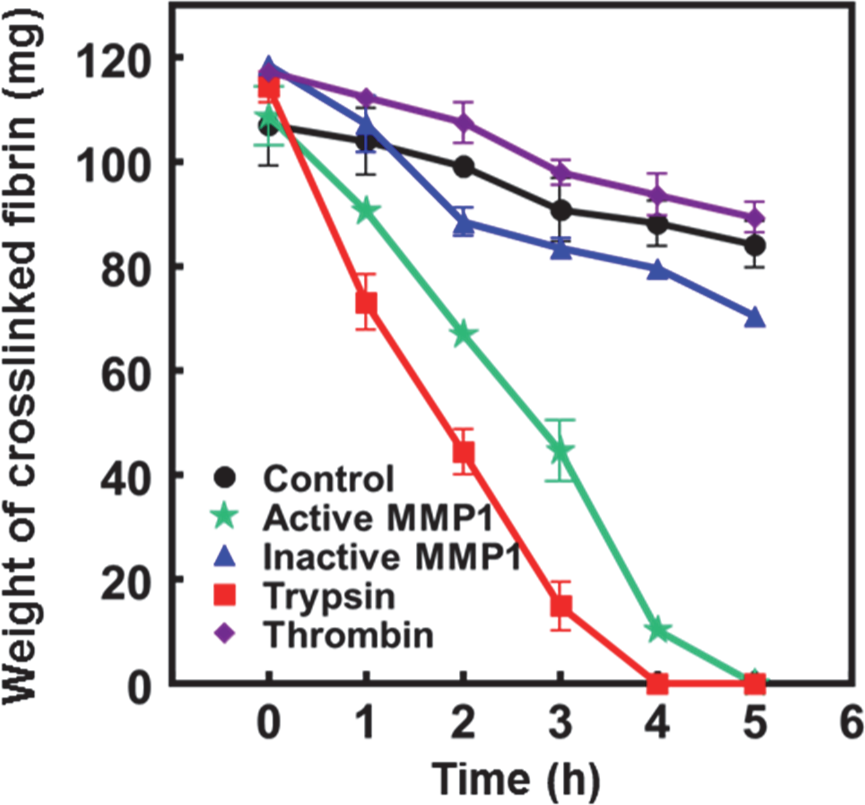
Weight-based fibrin degradation assay. Time-dependent weights of crosslinked fibrin after treatment with active MMP1, active site mutant of MMP1, trypsin, thrombin, and PBS buffer as the control.

### Weight-based degradation assay for water-insoluble fibrin

We used a weight-based degradation assay to quantify the activity of MMP1 because fibrin is water-insoluble and as such, solution-based biochemical assays are appropriate. We prepared fibrin with crosslinks by mixing 5 µg of human factor XIII, 20 units of thrombin (Cayman chemical, Cat# 13188), and 40 µL of 10 mM PBS (pH 7.4) in a 0.5 mL PCR tube at 22° C. We incubated the mixture for 10 min at 37° C without shaking. We added 100 µg of fibrinogen (Cayman chemical, Cat# 16088) and 20 µL of 5 mM CaCl_2_ to the mixture and incubated for an additional 15 min at 37° C without shaking. After incubation, the solution becomes turbid, indicating the formation of crosslinked fibrin. We prepared five reactions for crosslinked fibrin and added 1) 100 µL of 10 mM PBS (pH 7.4), 2) 100 µL of 1 mg/mL active MMP1, 3) 100 µL of 1 mg/mL active MMP1, 4) 100 µL of 1 mg/mL trypsin, and 5) 100 units of thrombin. We made the final volume of each reaction to be 200 µL by diluting with 10 mM PBS (pH 7.4). At 0 hr, we centrifuged the reactions at 10000 rpm for 10 min using a tabletop centrifuge, discarded the supernatant, and weighed the tubes. We subtracted the weights of the PCR tubes for each specific reaction before preparing the reactions. For five conditions, we measured weight of ∼120 mg. After weighing, we again added 1) 100 µL of 10 mM PBS (pH 7.4), 2) 100 µL of 1 mg/mL active MMP1, 3) 100 µL of 1 mg/mL active MMP1, 4) 100 µL of 1 mg/mL trypsin, and 5) 100 units of thrombin to a final volume of 200 µL by diluting with 10 mM PBS (pH 7.4). We incubated the reactions at 37° C without shaking. At 1 h, we centrifuged the tubes again, weighed, and subtracted the weight of empty tubes. We diluted again as above and incubated for another hour. We repeated the process at 0 h, 1 h, 2 h, 3 h, 4 h, and 5 h. We performed three biological repeats to calculate the mean and standard deviations. **Figure S5** shows that active MMP1 and trypsin degrade crosslinked fibrin at a similar rate of ∼30 mg/h. Active site mutant of MMP1, thrombin, and control show similar changes in weights at a rate of ∼7 mg/h. After subtracting the control, the rate of crosslinked fibrin degradation by MMP1 is ∼0.23 mg/h/μg. Note that the weight includes water and, as such, has a higher value.

**Figure S6.**
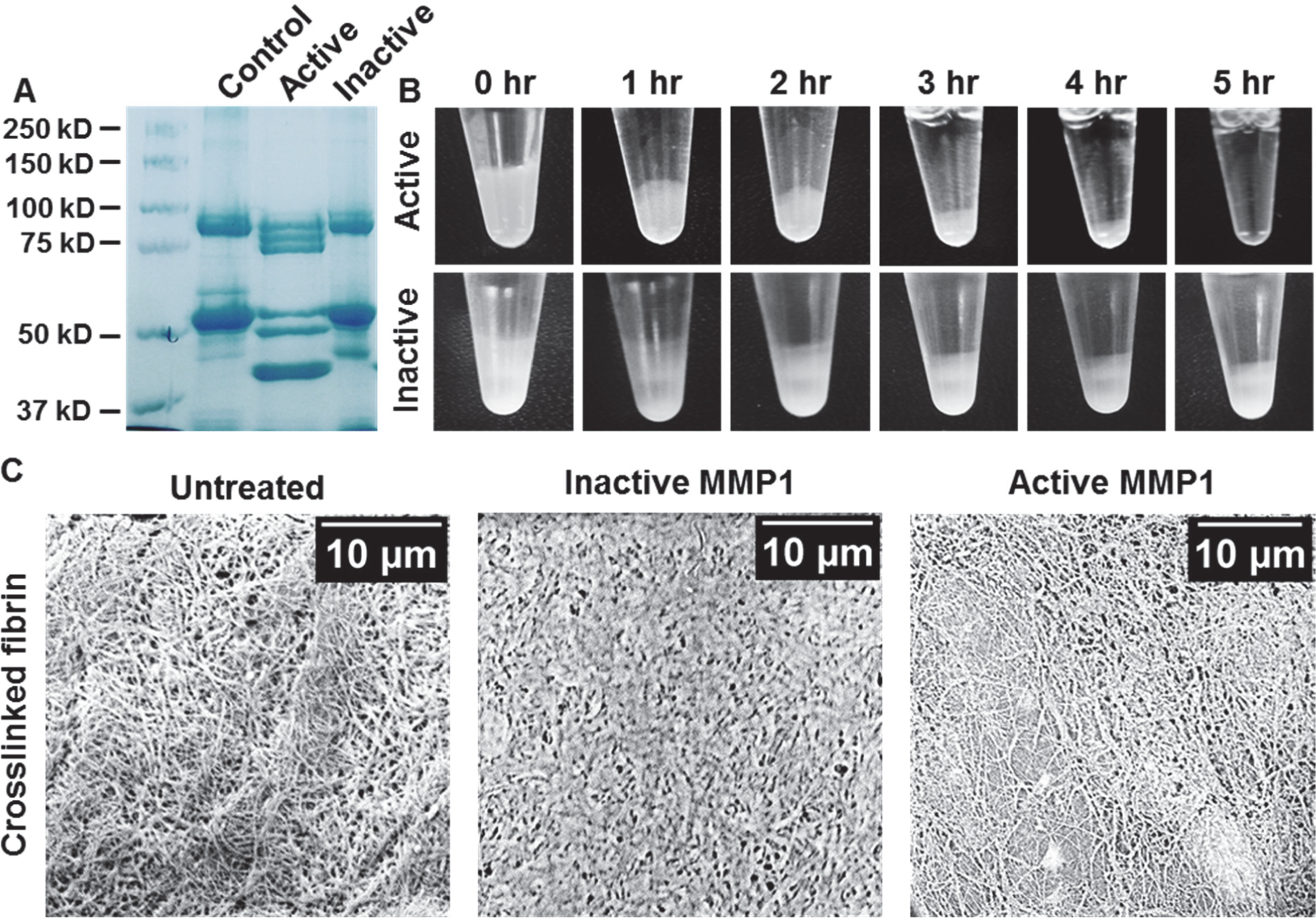
Fibrinolytic activity of MMP1 on crosslinked fibrin. (**A**) SDS PAGE of crosslinked fibrin with active MMP1 and active site mutant of MMP1. The control uses the protein buffer (50 mM Tris, 100 mM NaCl, pH 8.0). (**B**) 100 mg of wet crosslinked fibrin treated with 0.1 mg/mL active MMP1 and active site mutant of MMP1 at 37° C. (**C**) SEM images of crosslinked fibrin with and without MMP1 treatment.

### MMP1 activity on crosslinked fibrin

Water-soluble fibrinogen becomes water-insoluble crosslinked fibrin in the presence of factor XIII, thrombin, and CaCl_2_^10^. Thrombin converts fibrinogen into fibrin monomer by cleaving fibrinopeptide A and fibrinopeptide B^11,12^. Fibrin monomers self-assemble into protofibrils. The polymerization sites noncovalently attach to the D regions of two other fibrin(ogen) molecules. Fibrin monomers in each strand assemble end-to-end, whereas monomers across the strands arrange in a half-staggered overlap. The γ-chains of adjacent D regions in each strand is covalently attached by isopeptide bonds, which are formed due to the activity of factor XIIIa and appear as a ∼94 kDa band in SDS PAGE under reducing conditions (**Figure S6A**). Since factor XIIIa does not crosslink the β-chains^13,14^, the band at ∼52 kDa due to the β-chains remains unchanged in SDS PAGE of fibrinogen (**Figure S6A**) and crosslinked fibrin (**Figure S6A**). As shown in **Figure S6B**, active MMP1 degrades and dissolves water-insoluble crosslinked fibrin, but active site mutant of MMP1 does not dissolve crosslinked fibrin. **Figure S6C** shows the surface morphology of crosslinked fibrin imaged using SEM. Treatment with active MMP1 resulted in a more porous structure of crosslinked fibrin.

